# Journey to the center of the phage; revealing the ejectosome of *Pectobacterium* bacteriophage ΦM1

**DOI:** 10.1101/2024.02.19.577657

**Authors:** Alice-Roza Eruera, James Hodgkinson-Bean, Georgia L. Rutter, Francesca R. Hills, Rosheny Kumaran, Alexander J. M. Crowe, Nickhil Jadav, Klemens McJarrow-Keller, Fátima Jorge, Jaekyung Hyun, Hyejin Kim, Bumhan Ryu, Mihnea Bostina

## Abstract

Podophages that infect gram-negative bacteria, such as *Pectobacterium* pathogen ΦM1, encode tail assemblies too short to extend across the complex gram-negative cell wall. To overcome this, podophages encode a large protein complex (ejectosome) packaged inside the viral capsid and correspondingly ejected during infection to form a transient channel that spans the periplasmic space. Here we describe the ejectosome of bacteriophage ΦM1 to a resolution of 3.32 Å by single particle cryo-EM. The core consists of tetrameric and octameric ejection proteins which form a ∼1.5 MDa ejectosome that must transition through the ∼30 Å aperture created by the short tail nozzle assembly that acts as the conduit for the passage of DNA during infection. The ejectosome forms several grooves into which coils of genomic DNA are fit before the DNA sharply turns and goes down the tunnel and into the portal. In addition, we reconstructed the icosahedral capsid and hybrid tail apparatus to resolutions between 3.04 Å and 3.23 Å, and note an uncommon fold adopted by the dimerized decoration proteins which further emphasize the structural diversity of podophages. These reconstructions have allowed the generation of a complete atomic model of the ΦM1, uncovering two distinct decoration proteins and highlighting the exquisite structural diversity of tailed bacteriophages.

**Significance Statement:** This study resolves the cryo-EM structure of bacteriophage ΦM1, which possesses several unique and interesting structural elements, including a pair of distinct decoration proteins that are underreported in tailed DNA phages. Significantly, we also report the internal ‘ejectosome’ proteins of ΦM1, which are highly non-conserved with previously solved proteins to date and demonstrate the structural diversity of ejection proteins. The ejectosome reveals a DNA spooling phenomenon whereby the viral genome wraps around the ejectosome within the capsid, which has never been reported before. We provide a clear, step-by-step method for the technically challenging reconstruction of ejectosomes using standard, open-source software.

## Introduction

By current estimates, there are approximately 10^31^ bacteriophage particles in the biosphere, which equates to 10 times more bacteriophages on the planet than bacterial hosts (1, 2). Tailed bacteriophages of the order *Caudoviricites* are estimated to have emerged during the early precambrian period, making bacteriophages not only the most abundant biological agent on the planet, but also among the oldest (2, 3). Through such an extensive evolutionary history, phages have evolved exquisitely diverse and mosaic genomes, and are collectively capable of infecting essentially all known bacterial hosts (4, 5). Tailed bacteriophages are constrained to three major morphotypes based on tail morphology; the short non-contractile podophages, the long non-contractile siphophages, and the long contractile myophages. Despite significant conservation in structure, particle assembly and function amongst these morphotypes, mechanisms of host cell penetration and genomic ejection vary considerably (6).

Podophages belonging to the family *Autographiviridae*, otherwise known as the T7-like supergroup, possess non-contractile tails shorter than the width of the bacterial cell envelope (7-9). Some podophages translocate their genome into the host cytoplasm via a set of internal proteins known to form a protective channel that spans the entire width of the Gram-negative cell wall (10-12). Collectively, this internal complex is referred to as the internal core, or the ejectosome. The ejectosome is composed of an assembly of three proteins housed inside the capsid, coaxially arranged as rings mounted on the crown or barrel domain of the phage portal (13, 14). The ejectosome is expelled through the relatively narrow tail channel into the periplasmic space of the cell wall ahead of the genomic DNA. Within the periplasmic space, the proteins rapidly reassemble into a trans-envelope tunnel, subsequently acting as a conduit for the viral DNA to pass through the periplasmic space, preventing degradation by host nucleases or damage by the oxidizing periplasmic environment (8, 15). The T7 tetrameric ejection protein, gp16, encodes an LTase domain known to locally degrade peptidoglycan during this process, allowing complete penetration of the cell wall (8, 9). After genome delivery into the cytoplasm, the ejectosome rapidly disassembles and the cell wall is mended, preventing premature cell lysis (8, 14, 15).

Despite the recent publication of several high-resolution whole particle podophage structures known to encode ejectosomes (e.g. HRP29, P-SCSP1u, GP4, carin-1), only the *Escherichia coli* podophage T7 ejectosome has been successfully reconstructed (14-18). The reconstruction procedure is technically challenging and the lack of available reference models makes tertiary structure prediction tools like AlphaFold unreliable. Indeed, ejection proteins simply remain as annotated features in the sequenced genomes of many bacteriophages, or are occasionally identified as disordered densities in map slices through asymmetric reconstructions (15, 16, 18, 19).

In 1995, an investigation into new transducing phages for use as genetic tools in *Pectobacterium atrosepticum* lead to the discovery of bacteriophage ΦM1 (18, 19), which was later shown to generate mutants capable of escaping an abortive infection mechanism in the host induced by a Type III protein-RNA toxin-antitoxin system (20). The receptor remains unknown. Phage ΦM1 is grouped into genus “*Phimunavirus*”, of which members are referred to as “ΦM1-like” (21). Phage ΦM1 encodes a DNA genome approximately 44 kb long, organized into 52 putative genes. The host bacterium for ΦM1, *P. atrosepticum*, is a globalGram-negative potato pathogen found in both temperate and tropical regions, and is known to colonize the intercellular spaces of plant cells and degrade the cell wall, causing blackleg disease and aerial stem rot (22, 23). Recent studies have focused on identifying biocontrol agents against *P. atrosepticum*, including antimicrobial chemicals, zinc/silver nanoparticles, and antagonistic bacterial isolates (24-27). Comparatively little research has been done on the potential of phage-based biocontrols.

Using cryo-EM, we investigated the whole particle structure of bacteriophage ΦM1, identifying two decorations with relatively uncommon morphology, and a tail fiber/spike assembly with features similar to the recently published structure of Shigella phage HRP29 and phage T7. We describe an odd and unexpected capsid dimerization phenomenon, which we speculate represents an aberrant population of particles that form early in particle maturation. We pay particular focus to the structure of the ΦM1 ejectosome, which shows significant structural divergence relative to bacteriophage T7. We identify and describe a pattern of viral DNA spooling around the exterior of the ejectosome machinery. And finally, we provide a clear cryoSPARC-based workflow for reconstruction of podophage ejectosomes, and describe an intrinsic complication in the asymmetric reconstruction of podophages that has likely limited success in previous reconstruction attempts by others.

## Results

### Structural organization of bacteriophage ΦM1

Bacteriophage ΦM1 encloses the viral DNA within a highly decorated T=7 icosahedral capsid composed of HK97-like major capsid proteins (MCPs) and two similar, but distinct, accessory proteins (Figure 1a, 1b). The capsid houses a large ejectosome complex composed of tetrameric ejection proteins (TEP, gp48), octameric ejection proteins (OEP, gp49), and ejection protein 3 (EP3, gp50) arranged axially above the crown of the portal complex (gp35). An adaptor assembly (gp52) interfaces with the portal and six tail fibers (trimeric gp47), each of which appears composed of long helical bundles much like phage T7, as well as adapts trimers of spike protein (gp39) which may be homologous to those in phage P22, another podophage, and 9NA, a siphovirus (Fig. S11) (28). Gp39 was poorly ordered in our reconstructions and was not modelled. However, AlphaFold-2 multimer predicts a structure similar to phage HRP29 gp52. The short tail assembly has the closest similarity to bacteriophage HRP29 (16).

**Figure 1.**
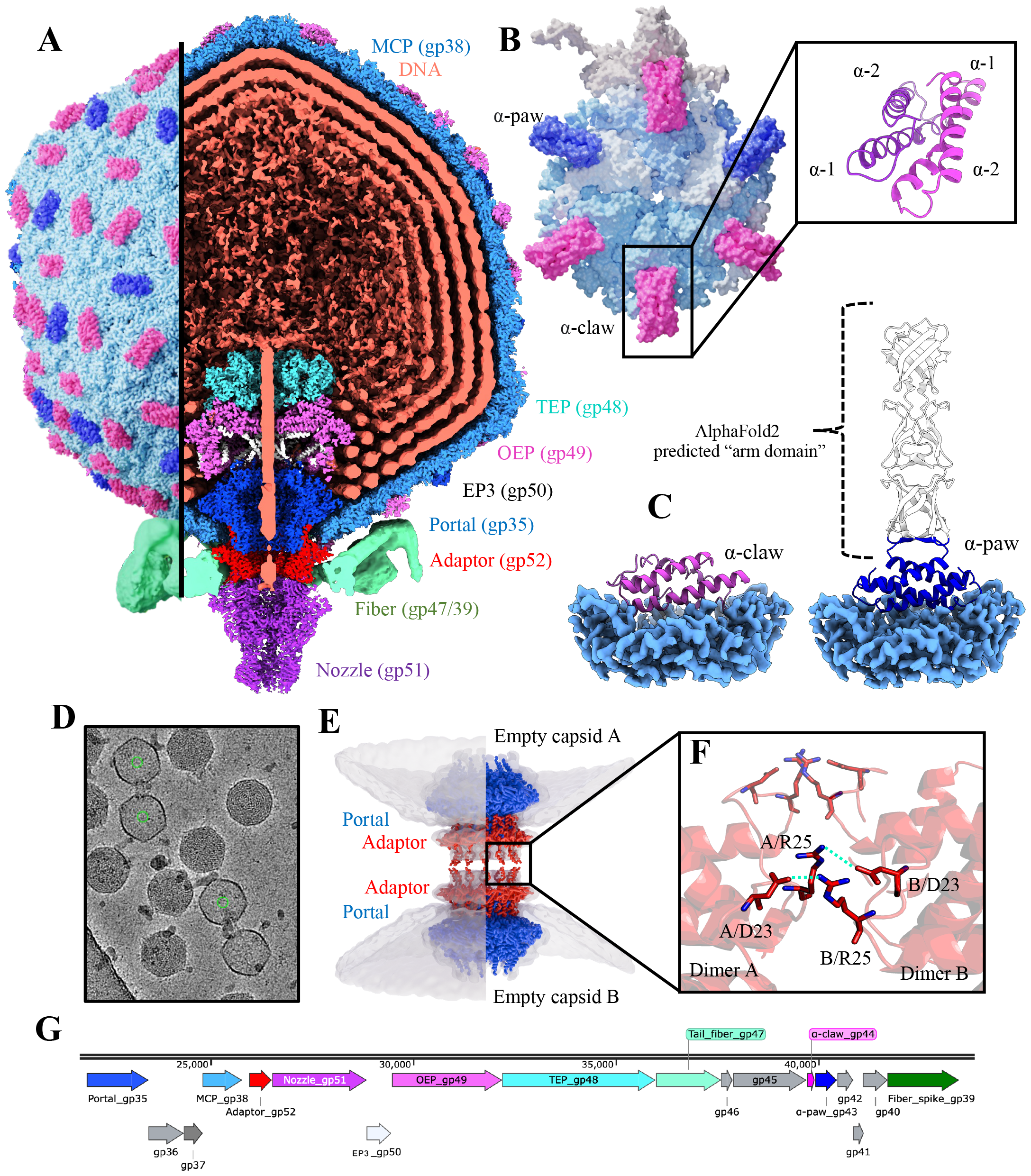
Overview of bacteriophage ΦM1. (*a*) Complete aligned composite reconstruction of the entire bacteriophage ΦM1 particle. (*b*) Asymmetric unit displayed in molecular surface representation. Homodimers of the α-paw protein (dark blue) and the α-claw protein (pink) decoration proteins that bind the 2-fold and quasi-2-fold axis in an alternating pattern dictated by pentamer proximity. MCPs that contact α-claw proteins are colored light blue. MCPs that contact α-paw proteins are colored white. The pentameric MCP is colored cream. Boxed on the right is a depiction of the α-claw motif which is conserved in both decoration proteins (*c*) *In situ* depiction of α-claw and α-paw proteins. The unmodelled α-paw arm domain predicted by AlphaFold2 multimer is shown in white. (*d*) representative cryo-EM micrograph depicting two capsid dimers observed on a single micrograph. (*e*) Models of the portal and adaptor rigid body fit into the capsid dimer reconstruction (9.3 Å). No DNA, ejectosome proteins or distal tail assemblies were resolved in the capsid dimer reconstruction, indicating the dimer forms at an adaptor-adaptor interface. (*f*) Opposite and opposed adaptor dodecamers showing a possible network of reciprocal salt bridges between Arg25 and Asp23 between opposing adaptor monomers. (*g*) Schematic of the ΦM1 structural module (generated in SnapGene V7.0.2).

Two decoration proteins, gp44 and gp43, are individually expressed off the viral genome and bind across MCPs at the the 2-fold and quasi-2 fold axes of symmetry via formation of homodimeric α-helical bundles, where each dimerization domain consisting of a helix-turn-helix motif (Figure 1c). We refer to these bundle structures as α-claw motif’s, as corresponding density bears resemblance to claw clips. The gp44 consists only of the α-claw motif, which we refer to as the α-claw protein. Gp43, however, also encodes a long helical fiber that we term the ‘forearm’ domain. We refer to gp43 as the α-paw protein. Whilst the α-claw is a small protein (∼6 kDa) and is primarily two α-helices, the α-paw is nearly three times the size (∼17 kDa), due to the long C-terminal forearm domain (Figure 1c). The forearm domain was disordered, likely due to high flexibility.

Empty capsids reconstructed from the data revealed a number of aberrant capsid-complex products which dimerize at the tail end into a structure reminiscent of gemini viruses (Figure 1d, 1e) (29). A low-resolution (∼10 Å) D12 reconstruction of a capsid dimer species showed density for the portal and adaptor complexes of the neck (figure 1d, 1e), but no density for the ejectosome, tail fibers or the nozzle. Inspection of the dimer interface between symmetrized adaptor models auto-fit into the map suggested the most likely explanation for this oligomerization may be potential reciprocal salt bridges between Arg25 and Asp23 of each chain of gp52 (Figure 1f). This hypothesis is tentative given the poor map resolution. Three species of dimers were observed in our data, empty-to-empty, empty-to-full and full-to-full capsids.

### The α-claw and α-paw decoration proteins

The ΦM1 particle is highly decorated with two accessory proteins, termed the α-claw, and the α-paw. Both proteins bind to 2-fold or quasi 2-fold axes, forming a clip-like structure between opposite and adjacent MCP monomers belonging to adjacent hexamers or pentamers (Figure 2a). The decorations are arranged such that the more common α-claw proteins are bound to all intercapsomeric interfaces, except those immediately flanking the hexamer-pentamer interface. As such, the ratio of α-claw:α-paw is 5:2 per asymmetric unit. The protein fold adopted by these decorations is under-reported, with only one other podophage described with a similar decoration protein fold (16).

**Figure 2.**
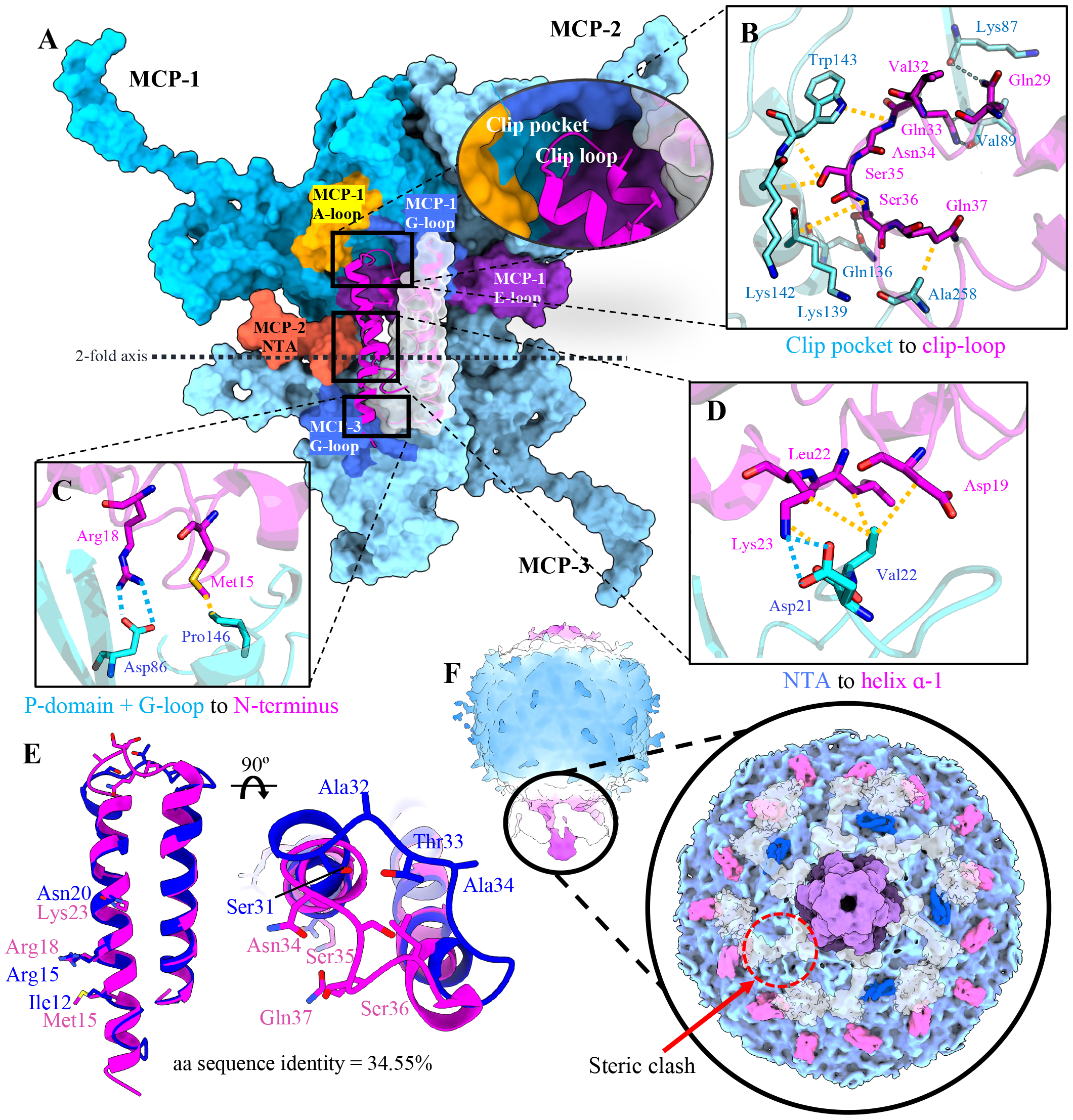
Binding of α-claw motif decoration proteins to the ΦM1 capsid. (*a*) ΦM1 α-claw-capsid surface interface. MCP monomers are labelled MCP 1-3 and are coloured different shades of blue to indicate monomer boundaries. A black dotted line shows the 2-fold axis. Major MCP features are coloured and labelled according to their identity. Circled is a close up of the conserved clip loop motif binding into the clip pocket, formed by surrounding MCP-1 A-loop (orange), G-loop (royal blue) and E-loop (dark orchid). (*b*) Interactions between the α-claw clip-loop and the MCP-1 clip pocket (hydrogen bonds are shown in blue, non-polar contacts are shown in yellow). (*c*) Interaction between MCP-3 P-domain and G-loop (blue) and the α-claw N-terminus (pink). (*d*) Interaction between the α-claw α-1 helix and the MCP-2 N-terminal arm. (*e*) Overlay of the α-claw and α-paw modelled regions, with key interacting residues highlighted to show conservation/divergence. The α-claw is shown in hot pink and the α-paw is shown in dark blue. (*f*) A low-pass filtered whole particle asymmetric reconstruction (left) and a segmented surface model (right) indicate the distribution of α-claw and α-paw around the unique portal vertex. MCP’s are shown in light blue, the nozzle is shown in purple, fiber/spike assemblies are shown in transparent white, the α-claw is shown in hot pink and the α-paw is shown in dark blue. Density for an α-paw is missing at a probable steric clash site where a fiber lays over the α-paw binding region.

Structure alignments of the conserved MCP-binding motif of each protein reveals a pruned RMSD of 0.865 Å (23 AA) and 8.17 Å across all 46 residues in the motif; additionally, amino acid sequence alignments of the same region showed 34.55% identity. Inspection of the dimerisation interface of both α-claw and α-paw homodimers using PISA analysis showed a strong hydrophobic contribution with only 2 hydrogen bonds predicted between α-paw monomers (−14.6 ΔG kcal/mol per α-paw dimer, -18.4 ΔG kcal/mol per α-claw dimer) (30). The major point of distinction appears to be in the ‘clip loop’ that connects each protein’s helix-turn-helix motif. In both cases, residues in this loop contribute many MCP contacts, and thus variation in each loop may contribute to the observed variation in binding pattern.

Both proteins make contacts with 3 MCP’s (referred to as MCP-1 to MCP-3 here). The α-claw first contacts MCP-1 via the clip-loop, which binds to the clip-pocket formed by the A-loop, Spine helix, E-loop and G-loop of an MCP on one side of the intercapsomeric interface. This is the largest α-claw-MCP interface, and appears mostly to involve hydrophobic interactions and hydrogen bonds (Figure 2b). A second interaction occurs with MCP-3 positioned across the intercapsomeric interface and is mediated by a putative salt bridge between Arg18 (α-claw) and Asp86 (MCP) (Figure 2c). A third interaction occurs with the N-terminal arm of MCP-2 forming a putative salt bridge between Asp21 (MCP) and Lys23 (α-claw), in addition to several hydrophobic contacts forming between Val22 (MCP), Lys23, Leu22, and Asp19 (α-claw) (Figure 2d). These monomeric interactions are then approximately reciprocated with the homodimeric partner in an antiparallel fashion. The α-paw protein also makes contacts with 3 MCP’s in the same manner, with some key differences. The α-paw clip-loop sits in a more shallow position in the clip-pocket and is predicted to make weaker interactions (Fig. S2). Additionally, the α-paw encodes an asparagine at position 20, which likely forms hydrogen bonds with Asp21 (MCP), in contrast to the putative salt bridge present at this location in the α-claw protein. Finally, a putative salt bridge is made between MCP Asp86 and Arg18 of the α-paw, or Arg15 of α-claw, respectively. Ultimately, PISA predicts a similar number of salt bridges and H-bonds between MCPs and each decoration protein, although the hydrophobic interface contribution is significantly greater in the α-claw (−1 ΔG kcal/mol per α-paw dimer, -8.6 ΔG kcal/mol per α-claw dimer).

As changes in the local capsid environment have been previously shown to modify the distribution of decoration proteins (31), we inspected how the unique portal vertex affects the distribution of both the α-claw and α-paw via asymmetric reconstructions. As might be expected, α-claw proteins that would normally bind the pentamer-hexamer interface were absent, due to the lack of reciprocating pentameric MCPs. The distribution of α-paw proteins immediately flanking the unique vertex were almost identical, with the exception that a single α-paw dimer was absent at a single interface (Figure 2f). This interface sits directly below one subunit of the tail fiber/spike complex, and we attribute this absence to steric hindrance.

### Organization of ΦM1 ejectosome

Similar to that of bacteriophage T7, the ΦM1 ejectosome is encoded as a three-protein mega complex composed of TEP, OEP and EP3 oligomers arranged coaxially within the phage capsid (Figure 3a, 3b, 3c). The EP3 octamer adapts the locally 8-fold symmetric interface of the ejectosome to the 12-fold symmetric crown of the portal by intercalating the EP3 helix α-2 between the crown domains of adjacent portal monomers, forming contacts with the C-terminal portal helix (Figure 3e, 3f). EP3 intercalates in a pattern in which two EP3 α-2 helices intercalate in the gaps between two adjacent portals, followed by an unoccupied gap. This results in 4 unoccupied portal crown domain gaps, and 8 filled with the α-2 helices of the EP3 octamer (Figure 3g).

**Figure 3.**
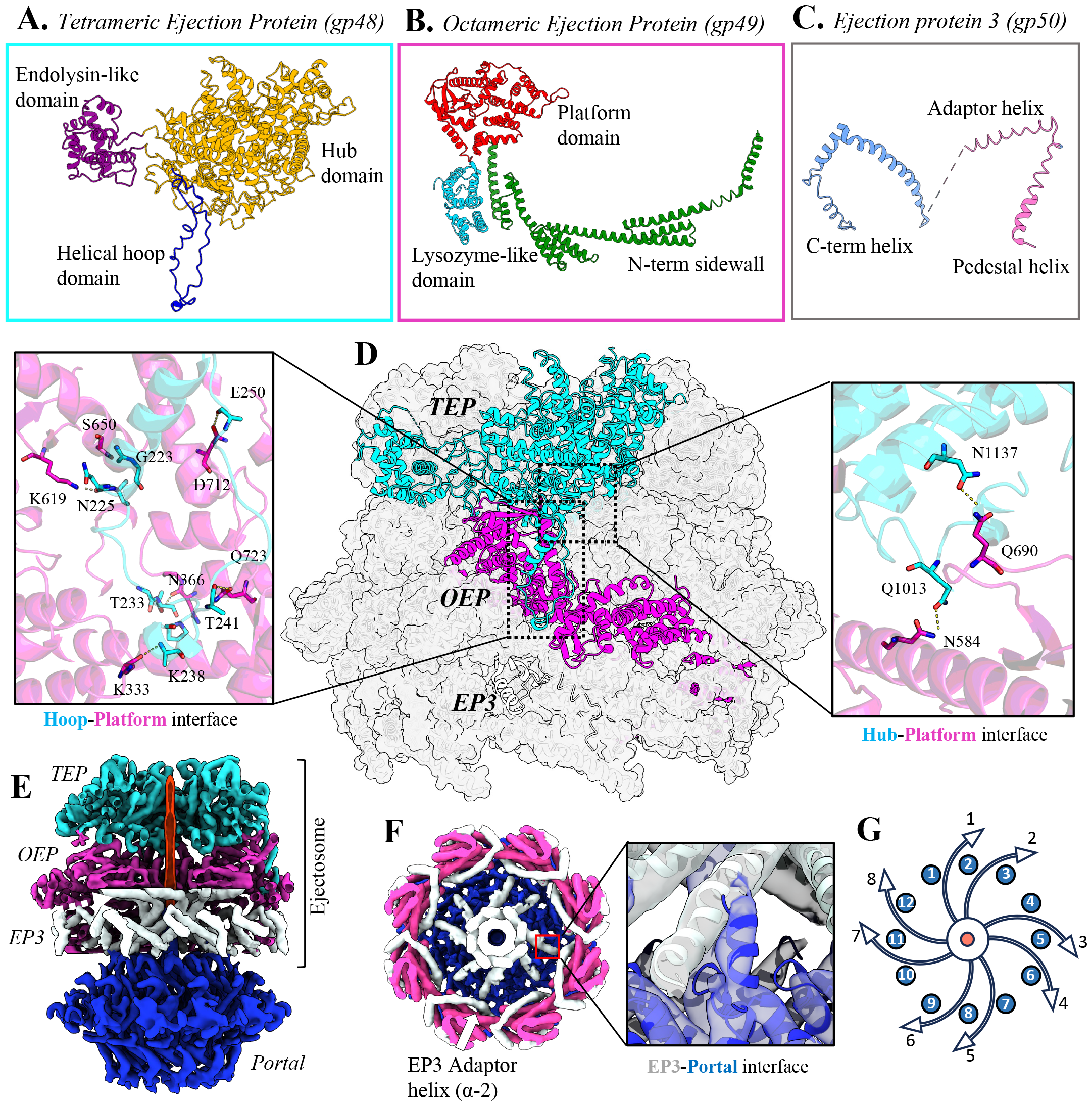
Architecture of ΦM1 ejectosome. Subdomains of tetrameric ejection protein (*a*), octameric ejection protein (*b*) and Ejection protein 3 (*c*) are labelled. Where a domain is sufficiently conserved with the homologue of bacteriophage T7, the naming convention of T7 has been preserved. (*d*) One chain of the TEP, OEP and EP3 is coloured, with symmetry mates of the respective assemblies displayed as silhouettes. Two interfaces of the TEP and OEP are displayed with relevant polar contacts shown between sidechains. (*e*) The consensus map of the ejectosome is displayed on top of the portal assembly with a strand of genomic DNA shown in red within the DNA channel. (*f*) A view of the top of the ejectosome looking down the DNA channel shows the alpha helixes of the EP3 adaptor, with a view of the helical contact interface with the crown of the portal complex. (*g*) A schematic of the axial arrangement at the EP3-portal interface. The EP3 α-2 helices (white arrows) adapt to the 12 helices of the portal crown. The central tube of DNA is represented in red.

The ΦM1 ejectosome differs significantly from that of T7 in structure, domain composition of the core proteins, interface interactions and potential functions (Fig. S3). Through a combination of BLASTP and DALI homology searches, putative enzymatic domains were identified in the ejectosome; specifically, a lysozyme-like domain within the OEP (Gly718-Ser903) and an endolysin/exotransglycosidase-like domain within the TEP (Asp268-Asp528) (32, 33). Viral lysozymes and exotransglycosidases can digest peptidoglycan, suggesting these domains may degrade local peptidoglycan surrounding the periplasmic channel once ejected from the phage particle during infection (34).

During our initial ejectosome reconstructions, portal, EP3 and OEP monomers appeared resolved, while TEP monomers were highly disordered. Through symmetry expansion and 3D classification, we were able to identify two 3D classes with TEP density; in each class, the EP3 and OEP were rotated 45° relative to the other class. A single class was chosen and was further refined with C4 symmetry imposed, however resulting reconstructions did not feature density corresponding to the portal. From this, we deduce that the ejectosome exists in multiple discrete particle species, in which portals cannot be aligned (Fig. S4AB). A similar phenomenon was previously described in which the portal of podophage SF6 exists in two discrete populations, separated by a 6° rotation about the Z axis (35). In our 3D classifications, we also identified two populations of particles in which ejectosome structures were aligned, but portal density was also rotated by ∼6° about the Z axis This suggests that the ΦM1 portal may also exist as two discrete particle populations (Fig. S4C).

### DNA-spooling around the ejectosome

During C4 reconstruction of the ejectosome, disordered density was observed spiralling the outer surface of the ejectosome within 10?15 Å of the OEP and TEP (Figure 4b, 4c). By applying a strong low-pass filter, we were able to connect these long densities to the density rings that surround the ejectosome and portal, often observed in asymmetric podophage structures and widely accepted as density corresponding to viral DNA (16-18). In each section of the C4 map, two strands of dsDNA follow two alternative routes around the ejectosome, before entering at equivalent positions rotated 90° relative to one another. The DNA spools upwards in a left-handed manner around the ejectosome towards the capsid center, before making a sharp arch bending downward to enter the central channel and extend into the tail. For each pair of strands, assumed to represent multiple local conformations of entry, one strand coils more tightly around the side of the ejectosome running along a groove formed by the lysozyme-like domains of the OEP, and one strand arches over the hub of the TEP.

**Figure 4.**
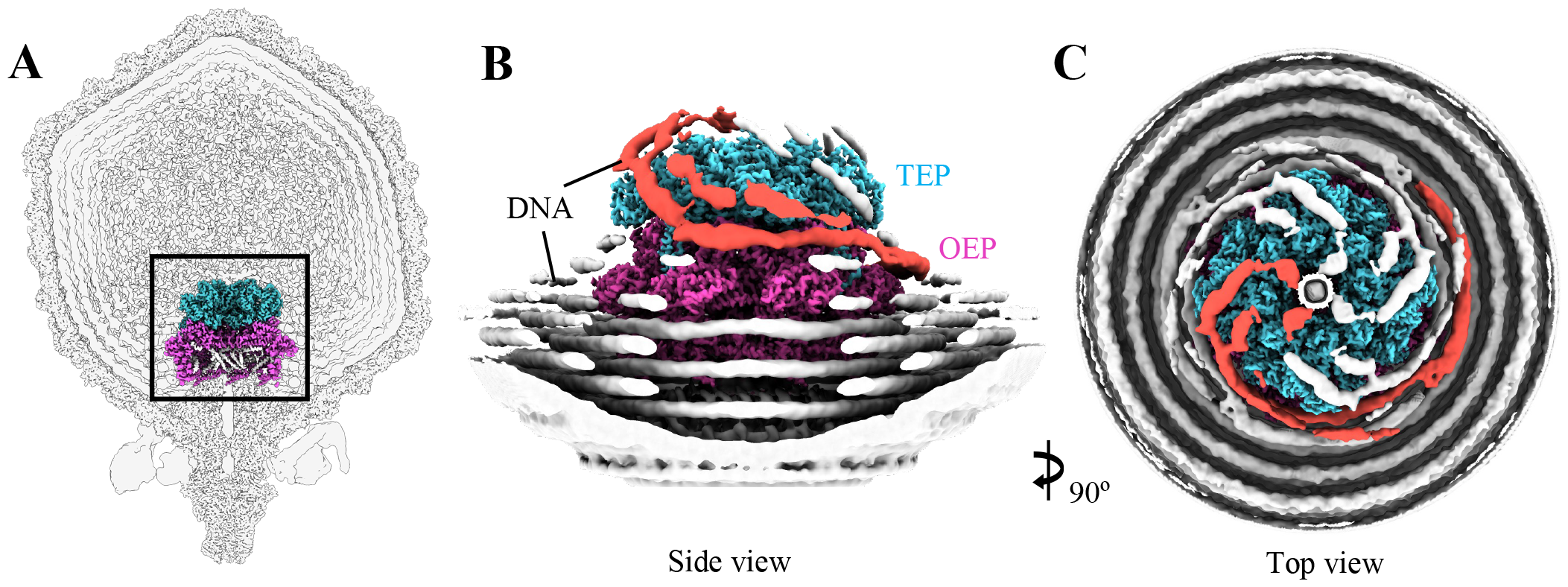
DNA spooling around the ΦM1 ejectosome. (*a*) A phage slice showing the relative location of the core proteins within the phage capsid. (*b*) A cropped view of the ejectosome (*TEP* cyan, *OEP* pink, EP3 not shown). DNA is shown in white with two strands, assumed to be rotational isomers of the same strand, coloured in red. (*c*) The same view is seen showing left-handed spooling of the DNA towards the central DNA channel.

Several attempts were made through focused 3D classification to isolate a single strand of DNA spooling the ejectosome. However, in all attempts, DNA still showed C4 symmetry. A single spool of DNA was measured from the point the strand separates from the lowest DNA ring to the top of the ejectosome, and was roughly 290?320 Å in length, corresponding to ∼85?94 base pairs of genetic information (1 base pair = 3.4 Å) (36).

## Discussion

ΦM1 can be thought of as an arrangement of protein oligomeric rings represented by C4, C5, C6, C8, and C12 symmetries. Such a wide array of symmetries creates several symmetry breaking interfaces which complicate the process of reconstructions with imposed symmetry. However, in this study, we discovered these boundaries also introduced difficulties in asymmetric reconstruction. The C12 portal complex that sits in the locally C5 MCP environment results in a symmetrically incompatible interface that is best characterized by C1 symmetry. However, if we consider the case of the portal/EP3 interface, and the OEP/TEP interface, multiple symmetrical positions become possible, complicating reconstruction procedures. Our workflow (symmetry expansion followed by 3D classification) permits the refinement of particle orientations such that all ejectosome components are aligned. However, the consequence of refining the alignment of ejectosome components was loss of the EP3/portal interface, and total loss of portal density. Further, during whole particle asymmetric reconstruction, capsids, nozzles, fibres and portals appear aligned, while the ejectosome showed visible but highly disordered density. Therefore, the TEP tetramer likely exists on top of the OEP octameric interface in two discrete rotamer species that cannot be aligned during asymmetric reconstruction (Fig. S4AB).

Analysis of other recent whole particle asymmetric podophage reconstructions (e.g. HRP29, P-SCSP1u, GP4, carin-1) suggests a similar phenomenon in which the ejectosome is visible in real space slices but was not reconstructed to high resolution. We hypothesize this might be caused by the improper sorting of whole particle rotamer species, resulting in misaligned internal proteins. By focusing only on the ejectosome, our reconstruction workflow allows ejectosome orientations to be refined such that ejectosome components are *correctly* aligned to each other, resulting in a high-resolution reconstruction with the lowest order of symmetry imposed (C4). Exploration of the extent to which rotamer species affect asymmetric reconstruction of bacteriophages represents a future area of investigation.

Capsid-like dimers were observed in our data and an empty-empty dimer was resolved to low-resolution, revealing a probable interface via a truncated tail apparatus. The empty-empty dimers may form via procapsid nucleation around a portal (without the ejectosome present), then dimerizing into the structure observed in our data (Fig. 1). If this is the case, the dimers may be an unintended aberrant product of capsid maturation. Alternatively, it is also possible the dimers form from two post-ejection phages which have lost the tail apparatus, later dimerizing at the adaptor interface, although there are at least twice as many polar contacts at interfaces of the nozzle and fibers to the adaptor (which would need to be lost) than there are for the adaptor to the portal (which is apparently retained) (Fig. S7). We note that a small population of whole empty particles (<60) with intact tails were also observed in our micrographs, indicating that genome ejection does not *always* result in the loss of the tail.

The α-claw and α-paw decoration proteins, although similar in the helix-turn-helix domain, may serve different functions during infection by virtue of their structures. Both decorations likely stabilize the capsid as they bind across MCPs, which is a common function for accessory proteins in large capsid viruses which can reach up to 60 atmospheric units of internal pressure once packaged (37). Given that the α-claw appears to form a stronger interface with MCPs, and doesn’t encode any additional fiber, we hypothesize that this protein is required to maintain capsid stability. In contrast, the α-paw likely contributes to stability to a lesser extent, but may mediate additional functions via the fibrous α-paw forearm domain, such as modulating host interactions via host recognition and adhesion. Similar functions have been observed in the decoration proteins of other DNA phages, e.g. phage N4 gp17, phi29 gp8.5, bacteriophage T5 pb10 and T4 HOC (37-40). Progressively truncating the α-paw protrusion and observing a decline in infectivity could confirm this hypothesis and could be pursued in future research.

DNA strands spooling around the ejectosome were observed in our low-pass filtered maps in two equivalent positions before turning sharply to enter the central channel and extend toward the tail. We suspect these different strands represent multiple conformations in which the final packaged DNA can interact with the ejectosome. Density is always observed to enter the ejectosome at approximately the same position, suggesting that some form of interaction constrains its path. From this, we speculate the ejectosome may play some role in the organization of DNA in ΦM1. Attempts to isolate a single DNA strand through 3D classification were unsuccessful, as in all cases, four densities corresponding to DNA could still be observed surrounding each TEP monomer. Whole particle asymmetric reconstructions may resolve this issue and shed further light on the nature of this interaction; however, given the intrinsic difficulties associated with true asymmetric reconstruction, a much larger dataset would be required for such an analysis.

## Conclusion

We describe an intrinsic complication in the asymmetric reconstruction of podophages that has likely limited success in previous attempts to reconstruct the ejectosome. The ejectosome proteins of ΦM1, despite clear structural homology, appear highly divergent both in structure and sequence when compared to the T7 ejectosome. In this study, we clearly describe a cryoSPARC-compatible reconstruction process to produce structures of the ejectosome of ΦM1 by single-particle cryo-EM. We describe genomic DNA spooling around the ejectosome in a left-handed manner before exiting the capsid and extending into the tail. In addition, two decorations bind the capsid across adjacent MCPs via a rarely observed structural motif, potentially stabilizing the capsid and engaging in host adhesion. Further, capsids or capsid-like products appear to dimerize at the connector interface, which may happen during an early stage in particle maturation or perhaps after genome ejection. The methods described in this study will support the efforts of future structural virologists in the reconstruction of symmetric and asymmetric elements of phages.

## Materials and Methods

Phage ΦM1 was purified and prepared for cryo-EM analysis as described in SI Appendix, SI Materials and Methods. Details of data collection, model building and analysis are also provided in the SI Appendix, SI Materials and Methods.

## Data Availability

The maps and models for the ΦM1 phage have been deposited in the Protein Data Bank (PDB) and Electron Microscopy Data Bank (EMDB), https://www.ebi.ac.uk/emdb/ with accession codes 8VB0 and EMD-43109 for the asymmetric unit, 8VB4 and EMD-43112 for the portal-adaptor assembly, 8VB2 and EMD-43110 for the ejectosome, 8VBX and EMD-43127 for the spike-nozzle assembly, respectively. The C8 ejectosome map and asymmetric whole particle map was deposited to the EMDB under EMD-43111 and EMD-43132, respectively.

## Acknowledgments

The authors would like to acknowledge the Institute for Basic Science in Daejeon, South Korea, and Sungkyunkwan University in Suwon, South Korea, for their contributions in sample preparation and data collection. The electron microscopy was supported by the National Research Foundation of Korea (NRF) grant funded by the Ministry of Science and ITC (grant number 2022R1A2C1005885). We would also like to acknowledge Joshua Voorkamp and Peter Higbee of the University of Otago for their wonderful technical IT support, and Max Wilkinson of MIT for advice on micrograph rescaling. We would like to thank Peter Fineran and his team for providing the initial phage sample for lysate preparation and purification. We acknowledge the staff at Otago Micro and Nano Imaging Center for their help in initial screening. The Otago Māori Early Career Researcher Fellowship contributed to the cost of transporting the specimen to South Korea. Finally, we would like to mihi to (acknowledge) the Kai Tahu peoples of Te Wai Pounamu, whose sovereign land provides the foundation of the University of Otago.

## References

1. G. F. Hatfull, R. W. Hendrix, Bacteriophages and their genomes. Curr Opin Virol 1, 298–303 (2011).

2. G. F. Hatfull, Bacteriophage genomics. Curr Opin Microbiol 11, 447–453 (2008).

3. H. W. Ackermann, Tailed bacteriophages: the order caudovirales. Adv Virus Res 51, 135–201 (1998).

4. D. M. Álvarez-Espejo, D. Rivera, A. I. Moreno-Switt, Bacteriophage-Host Interactions and Coevolution. Methods Mol Biol 2738, 231–243 (2024).

5. B. Ely, J. Lenski, T. Mohammadi, Structural and Genomic Diversity of Bacteriophages. Methods Mol Biol 2738, 3–16 (2024).

6. M. B. Dion, F. Oechslin, S. Moineau, Phage diversity, genomics and phylogeny. Nature Reviews Microbiology 18, 125–138 (2020).

7. S. Leptihn, J. Gottschalk, A. Kuhn, T7 ejectosome assembly: A story unfolds. Bacteriophage 6, e1128513 (2016).

8. N. A. Swanson, C. D. Hou, G. Cingolani, Viral Ejection Proteins: Mosaically Conserved, Conformational Gymnasts. Microorganisms 10 (2022).

9. S. R. Casjens, I. J. Molineux, Short noncontractile tail machines: adsorption and DNA delivery by podoviruses. Adv Exp Med Biol 726, 143–179 (2012).

10. I. J. Molineux, D. Panja, Popping the cork: mechanisms of phage genome ejection. Nat Rev Microbiol 11, 194–204 (2013).

11. I. J. Molineux, No syringes please, ejection of phage T7 DNA from the virion is enzyme driven. Mol Microbiol 40, 1–8 (2001).

12. B. Hu, W. Margolin, I. J. Molineux, J. Liu, The bacteriophage t7 virion undergoes extensive structural remodeling during infection. Science 339, 576–579 (2013).

13. X. Agirrezabala et al., Maturation of phage T7 involves structural modification of both shell and inner core components. Embo j 24, 3820–3829 (2005).

14. W. Chen et al., Structural changes in bacteriophage T7 upon receptor-induced genome ejection. Proc Natl Acad Sci U S A 118 (2021).

15. N. A. Swanson et al., Cryo-EM structure of the periplasmic tunnel of T7 DNA-ejectosome at 2.7 Å resolution. Mol Cell 81, 3145-3159.e3147 (2021).

16. S. Subramanian, S. M. Bergland Drarvik, K. R. Tinney, K. N. Parent, Cryo-EM structure of a Shigella podophage reveals a hybrid tail and novel decoration proteins. Structure 10.1016/j.str.2023.10.007 (2023).

17. L. Cai et al., Cryo-EM structure of cyanophage P-SCSP1u offers insights into DNA gating and evolution of T7-like viruses. Nature Communications 14, 6438 (2023).

18. A. d’Acapito et al., Structural Study of the Cobetia marina Bacteriophage 1 (Carin-1) by Cryo-EM. J Virol 97, e0024823 (2023).

19. L. Cai et al., Cryo-EM structure of cyanophage P-SCSP1u offers insights into DNA gating and evolution of T7-like viruses. Nat Commun 14, 6438 (2023).

20. T. R. Blower et al., Evolution of Pectobacterium Bacteriophage ΦM1 To Escape Two Bifunctional Type III Toxin-Antitoxin and Abortive Infection Systems through Mutations in a Single Viral Gene. Applied and Environmental Microbiology 83, e03229–03216 (2017).

21. C. Buttimer et al., Pectobacterium atrosepticum Phage vB_PatP_CB5: A Member of the Proposed Genus ‘Phimunavirus’. Viruses 10 (2018).

22. E. Monteagudo-Cascales et al., Study of NIT domain-containing chemoreceptors from two global phytopathogens and identification of NIT domains in eukaryotes. Mol Microbiol 119, 739–751 (2023).

23. I. K. Toth, Microbe Profile: Pectobacterium atrosepticum: an enemy at the door. Microbiology (Reading) 168 (2022).

24. E. Hamdy et al., Zinc Oxide Nanoparticles Biosynthesized by Eriobotrya japonica Leaf Extract: Characterization, Insecticidal and Antibacterial Properties. Plants (Basel) 12 (2023).

25. D. Elkobrosy et al., Nematocidal and Bactericidal Activities of Green Synthesized Silver Nanoparticles Mediated by Ficus sycomorus Leaf Extract. Life (Basel) 13 (2023).

26. J. Cigna et al., Efficacy of Soft-Rot Disease Biocontrol Agents in the Inhibition of Production Field Pathogen Isolates. Microorganisms 11 (2023).

27. J. E. Kang, S. Hwang, N. Yoo, B. S. Kim, E. H. Chung, A resveratrol oligomer, hopeaphenol suppresses virulence activity of Pectobacterium atrosepticum via the modulation of the master regulator, FlhDC. Front Microbiol 13, 999522 (2022).

28. D. Andres et al., Tail morphology controls DNA release in two Salmonella phages with one lipopolysaccharide receptor recognition system. Mol Microbiol 83, 1244–1253 (2012).

29. E. L. Hesketh et al., The 3.3 Å structure of a plant geminivirus using cryo-EM. Nature Communications 9, 2369 (2018).

30. E. Krissinel, K. Henrick, Inference of macromolecular assemblies from crystalline state. J Mol Biol 372, 774–797 (2007).

31. O. W. Bayfield et al., Structural atlas of a human gut crassvirus. Nature 617, 409–416 (2023).

32. L. Holm, C. Sander, Dali: a network tool for protein structure comparison. Trends in Biochemical Sciences 20, 478–480 (1995).

33. S. F. Altschul, W. Gish, W. Miller, E. W. Myers, D. J. Lipman, Basic local alignment search tool. J Mol Biol 215, 403–410 (1990).

34. C. Evrard, J. Fastrez, P. Soumillion, Histidine modification and mutagenesis point to the involvement of a large conformational change in the mechanism of action of phage lambda lysozyme. FEBS Lett 460, 442–446 (1999).

35. F. Li et al., High-resolution cryo-EM structure of the Shigella virus Sf6 genome delivery tail machine. Sci Adv 8, eadc9641 (2022).

36. M. Mandelkern, J. G. Elias, D. Eden, D. M. Crothers, The dimensions of DNA in solution. Journal of Molecular Biology 152, 153–161 (1981).

37. C. L. Dedeo, C. M. Teschke, A. T. Alexandrescu, Keeping It Together: Structures, Functions, and Applications of Viral Decoration Proteins. Viruses 12 (2020).

38. A. Fokine et al., Structure and Function of Hoc-A Novel Environment Sensing Device Encoded by T4 and Other Bacteriophages. Viruses 15 (2023).

39. K. H. Choi et al., Insight into DNA and Protein Transport in Double-Stranded DNA Viruses: The Structure of Bacteriophage N4. Journal of Molecular Biology 378, 726–736 (2008).

40. Y. Xiang, M. G. Rossmann, Structure of bacteriophage phi29 head fibers has a supercoiled triple repeating helix-turn-helix motif. Proc Natl Acad Sci U S A 108, 4806–4810 (2011).

