## Supplemental document for "Journey to the center of the phage; revealing the ejectosome of *Pectobacterium* bacteriophage ΦM1"

### Supporting Information Text

### Extended Methods

#### *Culturing of Pectobacterium atrosepticum and phage purification*

For lysate preparation, 6 mL of *P. atrosepticum* overnight culture was added to 200 mL low-NaCl LB (10 g/mL tryptone, 5 g/mL yeast extract, 5 g/mL NaCl), and was incubated for 1.5 hours at 25°C. To the culture, 400 mL of high-titre ΦM1 sample was added and incubated at 25°C for 17 hours. Crude lysates were centrifuged at 10,000 × g for 20 minutes (Thermo Fisher Lynx 6000, Fiberlite F12-6x500 LEX rotor) at 20°C and decanted into an autoclaved Schott bottle. Supernatant containing phage was filtered using a 0.22 µm filter. Sample was loaded into polypropylene thin walled 38.5 mL ultracentrifuge tubes (Beckman Coulter, ref: 326823) underlaid with 3 mL of 20% w/v sucrose and filled with phage buffer (10 mM Tris–HCl pH 7.4, 10 mM $MgSO_4$ and 0.01% w/v gelatin). Tubes were loaded into SW32 swing buckets. The sucrose cushion was spun at 50,000 × g for 1.5 hours at 20°C (Sorvall™ WX+ Thermo Fisher, Beckman SW 32 Ti rotor). Sample supernatant was decanted, and residual buffer was tapped out of tubes. All 6 pellets were resuspended in 500 mL of phage buffer overnight at 4°C. Pellets were pooled and diluted with phage buffer to a total volume of 22.5 mL.

Gradients were prepared using 4 × 1.5 mL of CsCl at densities of 1.33, 1.45, 1.6 and 1.7 g/cm$^3$ with the lightest density at the top and the heaviest density at the bottom, prepared in thin wall 17 mL polypropylene ultracentrifuge tubes (Beckman Coulter, ref: 337986). To both gradients, 11 mL of sample was loaded. Gradients were transferred into Beckman SW32.1 swing buckets and centrifuged at 100,000 × g for 4 hours at 20°C. Thick bands were visible post-centrifugation at the interface between 1.45 and 1.6 g/cm$^3$. Bands were carefully harvested followed by concentration and buffer exchanging into phage buffer using Amicon 100 kDa molecular weight cutoff 500 mL concentrators centrifuged at 14,000 × g at room temperature. Samples were stored at 4°C for later use.

#### *Negative-stain electron microscopy*

Specimens were assessed for sample quality and purity by negative-stain EM on a JEOL 1400 FLASH transmission electron microscope (TEM). Sample was loaded onto C-flat 300 nm mesh carbon coated grids followed by plasma discharge with negative charge using the GloQube discharge system (Quorum Technologies) for 30 seconds at 15 mA. Next, 5 µl of sample was applied to the carbon film and incubated for 60 seconds followed by negative staining with 5 µL of 1% phosphotungstic acid. Grids were air dried and imaged in the JEOL 1400 FLASH TEM operating at 120 kV, for sample quality, defined by intact particle appearance, good particle distribution and lack of obvious specimen contaminants. Of note, we observed almost all phage particles to have ejected their genomic material in the negatively stained images. This was in stark contrast with later cryo-EM imaging of the same phage sample which showed primarily full particles.

#### *Cryo-EM sample preparation and data collection*

Quantafoil R1.2/1.3 300 mesh grids were negatively glow discharged for 30 seconds at 45 mA and 3 µL of purified M1 phage was applied. Grids were plunge frozen using a FEI Mark IV Vitrobot (Thermo Fisher) with a blot force of 0, blot time of 3, 100% humidity and temperature of 4 degrees and wait time of 30 seconds. Data were collected using EPU software on two FEI Titan Krios G4 microscopes (Thermo Fisher), both operating at 300 kV with a K3 BioQuantum Gatan camera system. Exposures were collected for 12 seconds with an accumulated electron dose of 54 e/Å$^2$, fractionated into 50 frames. Exposures were collected with a range of defoci from -0.8 to -2.4 µm. The calibrated pixel size was 1.36 Å for one dataset and 1.40 Å for the other, and the lower pixel size data set was scaled for data merging purposes adapting the methods of Wilkinson *et al.* (1). A figure briefly detailing the scaling and merging method is available in Fig. S5.

#### *Data merging and processing*

Data were collected across multiple sites (Institute of Basic Science (IBS), Daejeon, and Sungkyunkwan University (SKKU), Suwon), each reporting slightly different pixel sizes (SKKU: nominal 1.40 Å, IBS: calibrated 1.36 Å). For data merging, the IBS dataset was rescaled to the larger pixel size of SKKU data to facilitate data merging for downstream processing (1). The reader is directed to Fig. S5 and Fig. S6 for a workflow diagram of the process. Both datasets were aligned, motion corrected and dose weighted using the Patch-Based Motion Correction function in cryoSPARC v4.2.1 (2). Icosahedral capsid reconstructions were then produced independently from a subset of both datasets using standard reconstruction methods outlined below, and were imported into UCSF ChimeraX for real space cross-correlation analysis between the two reconstructions using the fitMap tool (3). Peak cross correlation between the maps was observed at IBS: 1.394 Å, SKKU 1.400 Å. For the IBS dataset, micrographs were rescaled using this output to a final pixel size of 1.400 Å using Relion_Image_Handler (4). The micrographs were then imported into cryoSPARC and merged with the SKKU dataset. Micrograph stacks were then submitted to independent patch CTF correction. The combined micrograph stack contained 4,429 micrographs.

All downstream data processing was performed in version 4.2.1 of cryoSPARC. Briefly, global Contrast Transfer Functions (CTF) were fitted using the Patch-Based CTF Correction function and particles were picked using automated blob picking and subjected to iterative 2D classing until classes with clear secondary features were produced. Due to the modular nature of bacteriophages, downstream reconstruction workflow deviated significantly for different parts of the phage to best utilize available symmetry information.

***Capsid reconstruction***

For the capsid, multiple rounds of 2D classification yielded ~18,000 full capsid particles. Good particles were submitted to *ab initio* reconstruction without a symmetry group applied and the best class was selected for further refinement using the homogeneous refinement job with I2 symmetry imposed, followed by several rounds of non-uniform refinement in I2 symmetry with per particle defocus and negative Ewald sphere correction applied. Final DNA-full capsid maps were produced at a global resolution of 3.14 Å.

***Tail reconstruction***

For tail reconstructions, tail particles were identified by the following workflow. An early, strongly Fourier-cropped (4×) capsid reconstruction with imposed icosahedral symmetry was used to identify vertex coordinates spaced ~50 Å from the vertex surface using an in-house python script and UCSF ChimeraX. We note that distance from the vertex surface and box size appear to be important and sensitive variables in the success of this workflow and iterative optimization is strongly recommended. Particles aligned to this capsid reconstruction were symmetry expanded using I2 symmetry, and particle box centers were re-centered to the new vertex coordinates using a Volume Alignment Tools job in cryoSPARC. Particles were then extracted with a smaller Fourier cropped box size (360 pix → 180 pix), followed by removal of duplicate images using a 40 Å distance threshold. The particle stack (now ~12 × original size) was submitted to 3D classification without alignment. Vertex particles containing tails were pooled into one class approximately equal to the original stack size, which was then used as the unique vertex particle stack in downstream tail, fiber, and internal protein reconstructions.

Tail reconstruction was performed first by homogenous refinement with C6 symmetry imposed, followed by local CTF correction and non-uniform refinement. A final resolution of 3.23 Å was achieved for the full C6 symmetrised reconstruction. To resolve the C12 portal and adapter portions of the tail, particle centers were shifted 163 Å in Z, followed by reconstruction without alignment to produce an initial model with C12 components centered. This model was then refined with C12 symmetry using the local refinement job with a supplied focus mask, followed by re-estimation of local CTF and non-uniform refinement. The final full C12 symmetrised tail reconstruction resolution was 2.98 Å.

***Internal protein (ejectosome) reconstructions***

A full reconstruction protocol is available in Supplementary Method 1. Briefly, C12 tail coordinates described in the tail reconstruction section were re-centered to achieve a 165 Å shift in Z toward the capsid center, followed by re-extraction and reconstruction without alignments and a C4 local refinement with a large non-specific focus mask. Next, C12 symmetry expansion was performed followed by 3D classification to identify 3D classes with correctly aligned ejectosomes. Particles from one representative class were used for further local refinements in C1 symmetry. Through this method, C8 components corresponding to OEP and EP3 were recovered, however TEP appeared to be absent in reconstructions. To resolve this, symmetry expansion using C8 symmetry was performed followed by 3D classification, which yielded a complete ejectosome class for final local refinements. Duplicate particles were removed to effectively reverse symmetry expansion, resulting in a particle stack approximately equal to the initial stack size suitable for C4 reconstructions. This particle set was submitted to local CTF estimation and local refinement using a tighter focus mask as this method appears to yield the best results for proteins embedded in somewhat disordered DNA such as the portal and ejectosome. The final ejectosome reconstruction yielded a resolution of 3.32 Å. Note, the EP3 was modeled into the C8 ejectosome map but deposited in the C4 map.

***Model building and refinement***

Most preliminary models were produced in Alphafold2 using Google Colab (5) and fit into consensus maps, either segmented or whole, using UCSF ChimeraX (3) followed by manual refinement in COOT and ISOLDE (6, 7). In the case of the collar fiber trimer proteins, AlphaFold Multimer was used to produce the initial models using the Cosmic$^2$ server (8). For the TEP, which was particularly challenging given the uniqueness of the structure and many disordered loops within the map, an initial model was generated in AlphaFold2 but required additional efforts as only a small domain fit in the segmented map. The homologous protein to TEP from bacteriophage T7, which possesses a small conserved domain, was fit into the TEP map as best as practicable and the two models were then merged via Swiss Model (https://swissmodel.expasy.org) to create a better fitting model for the density (9). Most of this model then refined into the map as usual, although the first 300 N-terminal residues required modeling individually. Geometries were corrected in COOT and ISOLDE, and models were symmetrized to fit into the final consensus map. Model validation was performed by Molprobity score (https://molprobity.biochem.duke.edu/) (10).

**Supplementary method 1. CryoSPARC-based reconstruction protocol for resolving ejectosomes from podophages.**

*Prerequisites*
Phage tail particle coordinates (e.g., a particle stack of tail picks)
UCSF ChimeraX v2 or better
Any version of cryoSPARC up to at least version 4 (most current version at time of publication)

*Foreword*
This protocol is intended for use in cryoSPARC but the principles should be broadly applicable for any cryo-EM processing software of choice. This protocol first assumes a DNA-full phage tail class has already been obtained by either Homogeneous Refinement or Non-Uniform refinement. Unless otherwise specified, job types are run with default parameters (although the investigator is encouraged to change parameters important to their specimen of interest).

*Protocol*
To resolve an ejectosome, particle coordinates need to first be realigned so that the *center coordinate* of the particles are over the general location of the *ejectosome* (and not in the center of the *tail*). After this, reconstruction methods are relatively standard.

1. Obtain a reconstruction of the tail apparatus via Homogeneous Refinement or Non-Uniform Refinement.
    i. Note: It is important that symmetry is imposed in this reconstruction. Symmetry imposed may be either C6 or C12; the important criteria is that a high-quality portal structure is present in the tail reconstruction.
2. Import the particle coordinates from either refinement job into the Volume Realignment Tools job in cryoSPARC.
3. Shift the particle box along the Z axis so that the center of the particle box is roughly centered over the expected location of the ejectosome (i.e. within the capsid, above the portal).
    i. Note: The distance to shift will depend on the specimen being reconstructed and may require iterative optimization. In the case of bacteriophage ΦM1, shifting 165 Å along the Z axis worked well. We recommend starting with 165 Å as a preliminary value.
4. Import the new, realigned particle coordinates from the Volume Realignment Tools job into an Extract Particles job. Extract using appropriate parameters for your specimen.
    i. Note: The box size will need to be large enough to contain the ejectosome but small enough that minimal DNA is included within the box. DNA is a high-contrast, poorly ordered molecule and will overpower the weight of the ejectosome when the particles are aligned in downstream steps. Running a few Extract Particles jobs with progressively smaller box sizes may be beneficial, until an appropriate size is found which contains the ejectosome but minimal DNA.)
5. Submit the particle stack from the Extract Particles job (step 4) to a C12 Symmetry Expansion job.
    i. Note: At this point, masked local reconstructions of the ejectosome region will likely yield poor results, as tails will be aligned such that the ejectosome is rotated in several symmetry-related positions relative to the portal, resulting in disordered density. Symmetry Expansion allows re-insertion of each particle image into the Fourier shell in all symmetry-related positions. Then, by focused 3D Classification, particle images that have ejectosomes aligned in the same orientation may be pooled for further refinements.
6. Import the output from the Symmetry Expansion job into a 3D Classification job and run it with default parameters.

   i. We strongly recommend the use of a soft focus mask centered on the ejectosome region. A general blob shaped mask worked well in the case of ΦM1. (For soft mask generation, the investigator is directed to the cryoSPARC mask guide; [https://guide.cryosparc.com/processing-data/tutorials-and-case-studies/mask-selection-and-generation-in-ucsf-chimera](https://guide.cryosparc.com/processing-data/tutorials-and-case-studies/mask-selection-and-generation-in-ucsf-chimera).)
   ii. If <1,000,000 particles are available, we recommend selecting for no more than 3-5 classes. If >1,000,000 particles are available, expanding the search range through more classes may be beneficial. As there are 3 symmetry-related positions available for the ejectosome, normally 3 good classes should emerge, each ~4 times the size of the original particle stack.

**7.** Inspect the output volumes from the 3D Classification job. A 'good' class should contain some helices corresponding to the octahedral components near the portal.

**8.** Run a Remove Duplicate Particles job on any of the 'good' classes. The particle stack size should now be roughly equal to the initial particle stack size.

**9.** Submit the particles from the Remove Duplicate Particles job to a C8 Symmetry Expansion job.

**10.** Import the particle stack from the C8 Symmetry Expansion job to a 3D Classification job, with the same number of classes as in step 8 (again, a focus mask is recommended).

   i. Note: Here we expect *two* good classes, as tetrameric ejectosome components may sit in two symmetry-related positions on top of the octameric region.

**11.** Inspect the output volumes from the 3D Classification job.

**12.** Either good class should contain a fully aligned ejectosome. This volume can be used as an initial model, and/or as a template for local mask generation, in focused local refinement (after running a Remove Duplicates job). We found masked focused local refinement (without particle subtraction) worked best, presumably as DNA is excluded during refinement.

**Table S1. Sample preparation and data collection statistics.**

| | | **Sungkyunkwan University data set** | **Institute for Basic Sciences data set** |
|---|---|---|---|
| Grid preparation | EM-Grid | Quantifoil R1.2/1.3 Cu300 | Quantifoil R1.2/1.3 Cu300 |
| | Glow discharge (GD) | Negative | Negative |
| | GD current | 15 mA | 15 mA |
| | GD time | 30 seconds | 30 seconds |
| | GD pressure | 0.39 mbar | 0.39 mbar |
| Vitrification | Loading volume | 4 µL | 4 µL |
| | Loading side | Carbon | Carbon |
| | Blot time | 5 seconds | 3 seconds |
| | Blot force | 0 | 0 |
| | Temp/humidity | 4°/100% | 4°/90% |
| | Wait time | 0 seconds | 10 seconds |
| Microscope | Model | Krios G4 | TFS Krios G4 |
| | Spherical Aberration; Cs [mm] | 2.7 | 2.7 |
| | Dose rate [e/pix/s] | 8 | 8.3 |
| | Pixel value [Å/pix] | 1.4 | 1.36 |
| | Nominal Magnification | 64,000 X | 64,000 X |
| | Calibrated Magnification | 36,765 X | 36,765 X |
| | Exposure time [sec] | 12.25 | 12.02 |
| | # of Fractions | 50 | 50 |
| | Condenser lens aperture [µm] | 70 | 70 |
| | Objective lens aperture [µm] | 100 | 100 |
| | Defocus range (step size) | -0.8, -1.0, -1.2, -1.5, -1.8, -2.1, -2.4 | -0.8, -1.0, -1.2, -1.5, -1.8, -2.1, -2.4 |
| | Total dose [e/Å$^2$] | 53.38 | 53.9 |
| | Dose per fraction [e/Å$^2$/Frac.] | 1.07 | 1.07 |
| | AFIS (Fast Mode) | None | None |
| | Alpha Tilt [degrees] | 0 | 0 |

| | | | |
|---|---|---|---|
| Data | Model | Gatan K3 BioContinuum | Gatan K3 BioContinuum |
| | Energy slit width | 20 eV | 20 eV |
| | Acquisition mode | EC | EC |
| | Correlated double sampling (CDS) mode | Used | Used |
| | Movie format | Tiff(LZW) | Tiff(LZW) |
| | Gain normalization | No | No |

**Table S2. Reconstructed protein components of bacteriophage ΦM1.** The colours are consistent with figure elements and representations shown in this paper. All reconstructed proteins are listed. All molecular weights (MW) were estimated in ExPasy based on the amino acid sequence (https://web.expasy.org/protparam/). In the case of gp43 and gp44, the assembly is a dimer. The NCBI Taxonomy ID for bacteriophage ΦM1 is 1211386 and GenBank accession number YP_009591954.1. A single phage particle is composed of 23.8 Mda of protein.

| Protein name | Gp ID | Symmetry group | MW (monomer) | MW (assembly) | Monomers per phage (n = 909) |
|---|---|---|---|---|---|
| Major capsid protein | gp38 | I,222r | 36.4 kDa | 15 MDa | 415 |
| α-paw decoration | gp43 | N/A | 17 kDa | 34 kDa (per dimer) | 118 |
| α-claw decoration | gp44 | N/A | 6 kDa | 12 kDa (per dimer) | 290 |
| Tetrameric ejection protein (TEP) | gp48 | C4 | 135.2 kDa | 540.8 kDa | 4 |
| Octameric ejection protein (OEP) | gp49 | C8 | 97.9 kDa | 783.2 kDa | 8 |
| Ejection protein 3 | gp50 | C8 | 21.5 kDa | 172 kDa | 8 |
| Portal protein | gp35 | C12 | 55.6 kDa | 667.2 kDa | 12 |
| Adaptor protein | gp52 | C8 | 21.1 kDa | 253.2 kDa | 12 |
| Fiber | gp47 | C6 | 55.8 kDa | 167.4 kDa (trimer) | 18 |
| Nozzle protein | gp51 | C6 | 84.4 kDa | 506.4 kDa | 6 |

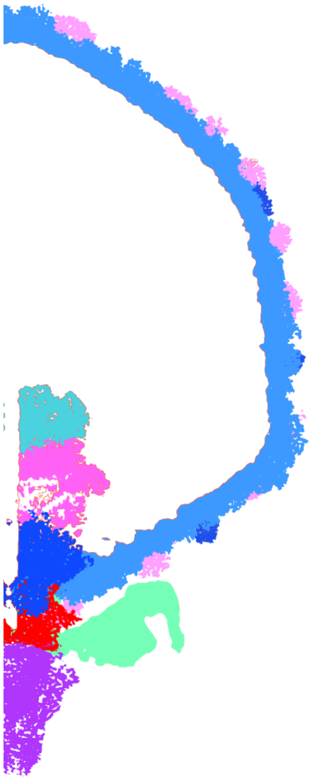

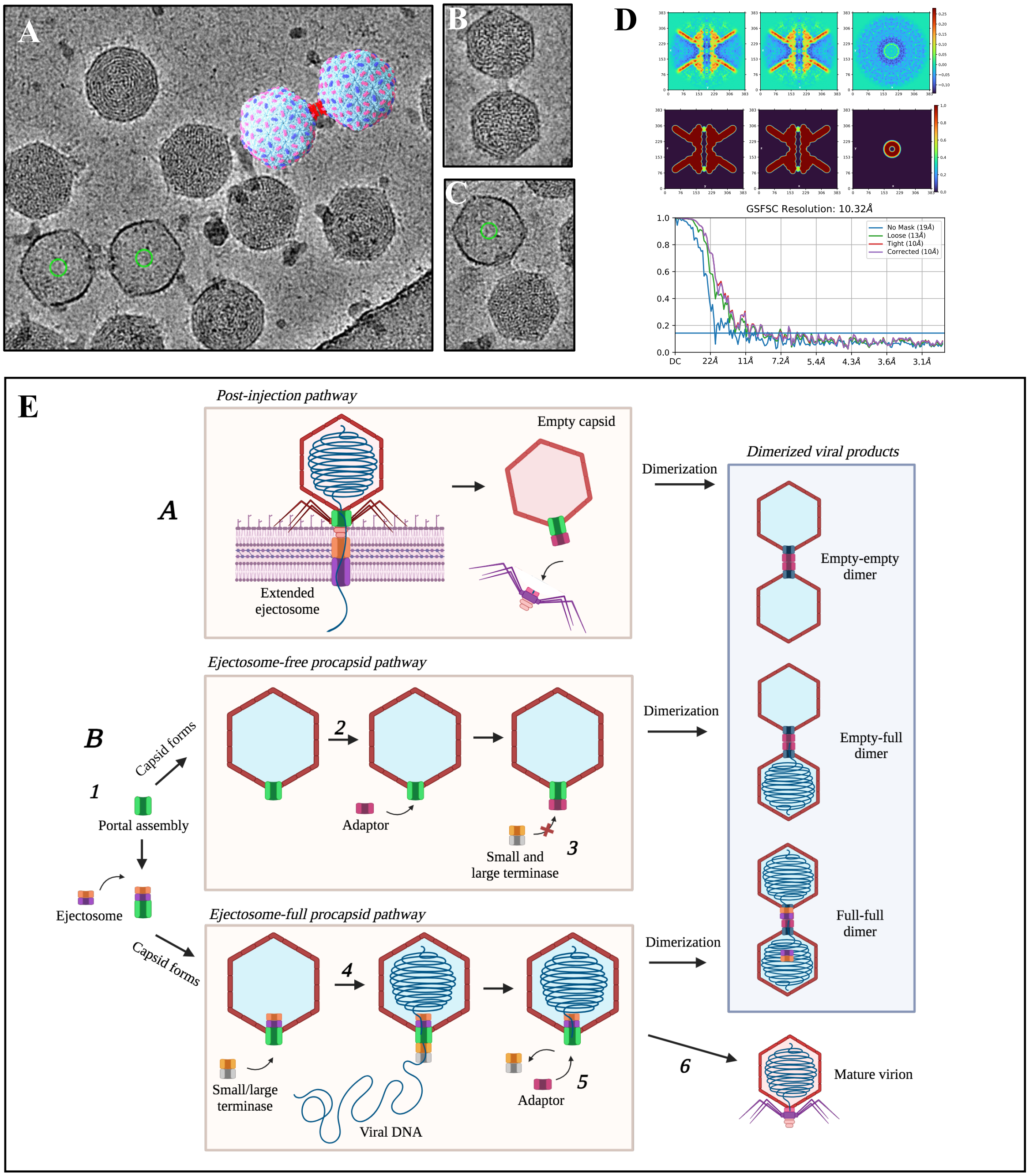


**Fig. S1. Immature dimerized ΦM1 products.** (*a*) Cryo-EM micrographs of three classes of immature capsid-like dimers. An artistic 3D render of the capsid dimer is placed over a dimer in the micrograph. The green circle indicates the center of a particle pick. Dimers can also be composed of two full (*b*) or empty-to-full capsids (*c*), but neither could not be resolved due to low particle counts (<60 particles each). Each class of capsid dimers (empty-empty, empty-full and full-full) were manually counted in the 3500+ micrographs of the merged data set. The empty-empty dimers were the most common product observed in these data. (*d*) Real-space slices and the gold-standard Fourier shell correlation plots for the empty-empty structure produced in cryoSPARC v4.2.1 (*e*) Two hypotheses can be proposed for the formation of the viral dimer products. Hypothesis A proposes that the ejectosome, DNA and tail assembly are lost after injection and the empty capsids dimerize into the empty-empty or empty-full products. Hypothesis B proposes the portal assembly (*1*), either with or without an ejectosome assembly associated to the portal crown, nucleates the formation of the viral procapsid, subsequently producing two capsid products (ejectosome-full and ejectosome-free). In the ejectosome-free pathway (*top*), the procapsid associates with an adaptor (*2*), at which point DNA packaging is no longer possible as the terminase cannot contact the portal assembly (*3*). The ejectosome-free procapsids dimerize with each other or with a full procapsid at the adaptor-adaptor interface. In the ejectosome-full pathway (*bottom*), the terminase associates with the portal and packages the viral genomic DNA in an ATP-dependent manner (*4*). A dissociation signal is recognized by the terminase which causes the terminase to dissociate from the portal, and an adaptor then associates (*5*). The procapsid can then dimerize with an empty or full procapsid (to form an empty-full or full-full gemini) or can associate with the rest of the tail assembly to form a mature virion (*6*).

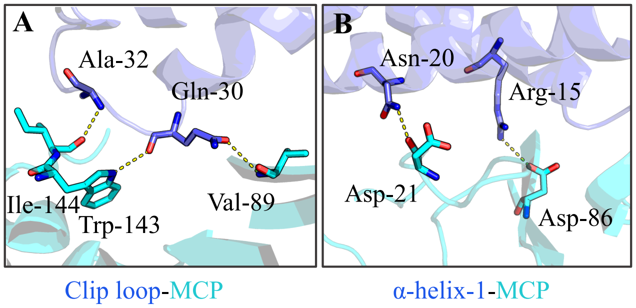


**Fig. S2. ΦM1 α-paw.** Reciprocal interface interactions which occur between the clip loop (a) and α-helix-1 of the α-paw and the major capsid protein.

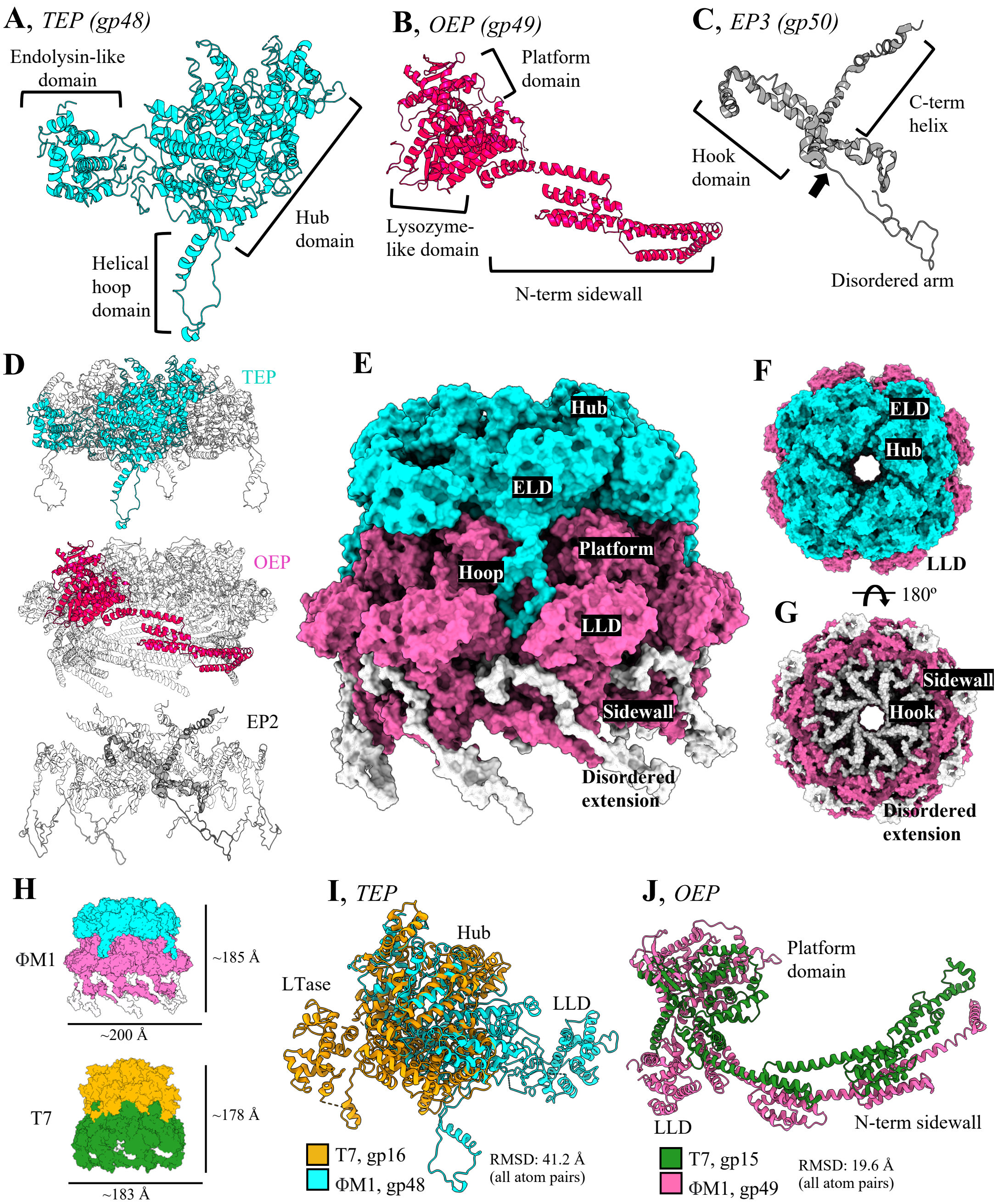


**Fig. S3. Subdomains of the ejectosome.** (*a*) Tetrameric ejection protein, (*b*) octameric ejection protein (*c*) and EP3 are labelled. Density in the EP3 cryo-EM map terminates near the arrow. The proposed structure of the N-terminal disordered extension is derived from AlphaFold2 and is preserved in this figure to give the reader an indication of the true length of the ejectosome assembly. (*d*) One chain of the TEP, OEP and EP3 is coloured, with symmetry mates of the respective assemblies displayed as silhouettes. (*e*) Molecular surface of the ejectosome is displayed from the side with domains labelled. (f) A view of the top of the ejectosome looking down the DNA channel shows the grooves in the hub which seat the DNA as it enters the tail assembly. (*g*) A bottom view shows the C8 arrangement of the EP3 hooks which interfaces with the top of the portal assembly (not shown). (*h*) A size comparison of ΦM1 ejectosome and T7 (7EYB) shows

ΦM1 is both wider and taller. A structural overlay of the TEP (*i*) from bacteriophage T7 (PDB: 7EYB) and the OEP (*j*) show similar general architecture but little conservation. Note, the EP3 model from T7 was too short to produce a meaningful overlay. RMSD was calculated in UCSF ChimeraX. A DALI structure database search was performed for the ELD, hub, platform, sidewall and LLD to probe for similar protein structures. DALI returned similar matches only for the endolysin-like domain of TEP and the lysozyme-like domain of OEP, matching with endolysin and lysozyme, respectively.

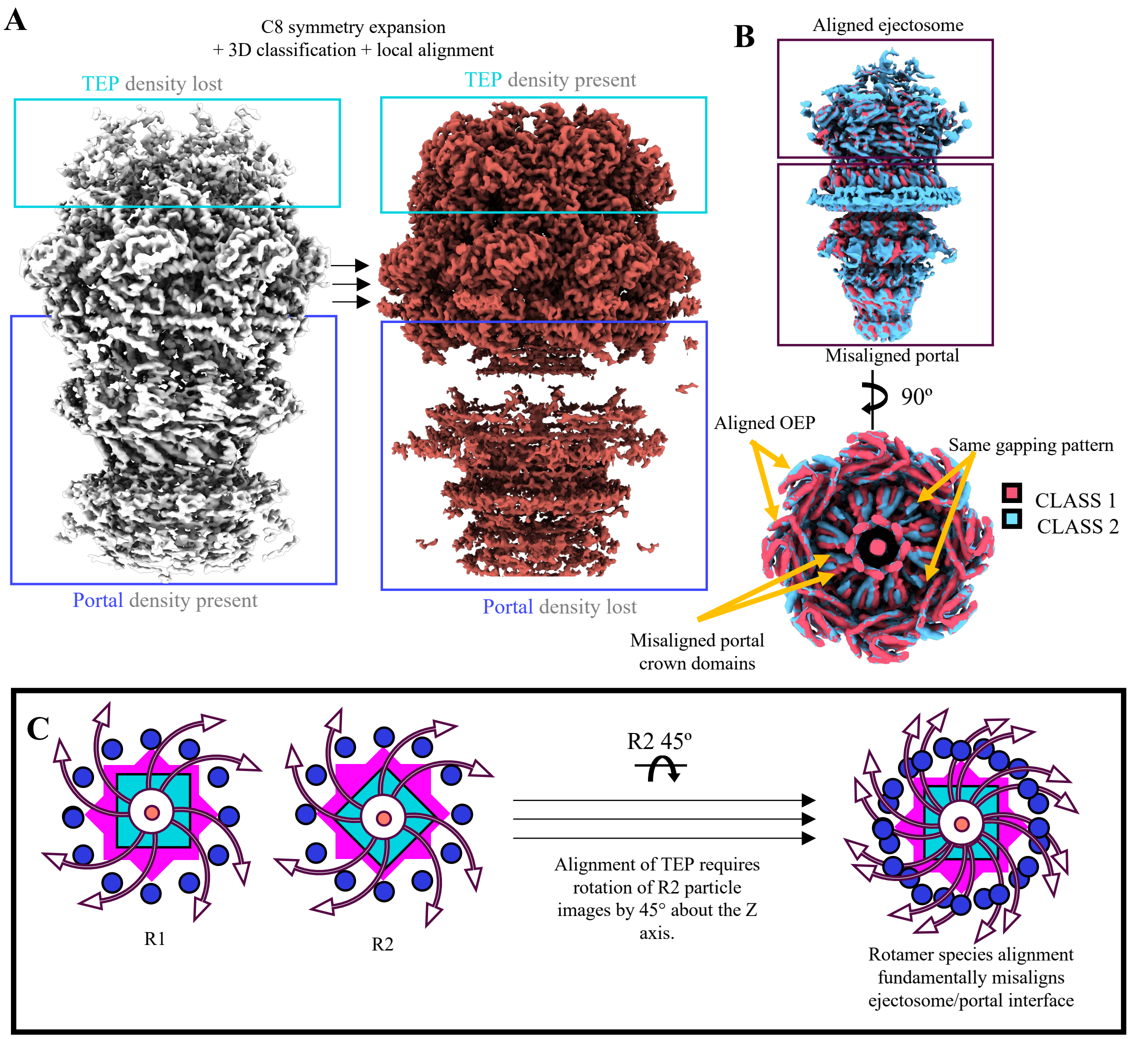


**Fig. S4. Evidence for the presence of ejectosome rotamer species.** (*a*) Ejectosome orientations were established by first aligning portal images, followed by a box shift in Z, and C12 symmetry expansion + 3D classification to find particle orientations with the portal correctly aligned to the ejectosome. TEP density was missing as shown in the cyan box. Portal density was visible but smeared; this is suspected to occur due to the presence of portal rotamer species, in which the portal rotates into two positions separated by 6° about the Z axis (below). C8 symmetry expansion and 3D classification refined particle orientations such that density corresponding to the TEP became visible at the expense of density corresponding to the portal. (*b*) Two representative 3D classes that show portals rotated by 6° relative to the ejectosome. (c) Theoretical depiction of how the presence of TEP rotamer species fundamentally prevents alignment of the portal and full ejectosome using a complete particle stack. Two rotamer species are depicted as R1 and R2; each species is identical except with the exception that TEP tetramers are rotated 45° about the Z axis relative to each other.

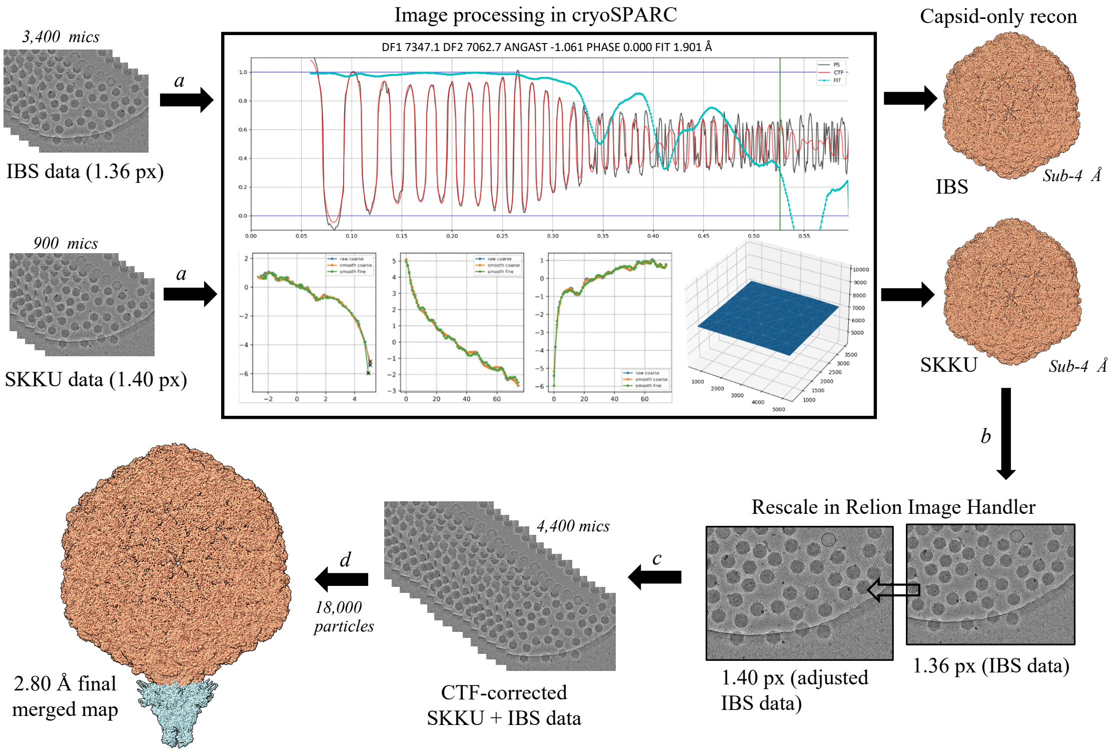


**Fig. S5. Data merging.** Two datasets with different pixel sizes (IBS, 1.36; SKKU, 1.40) were image processed (*a*) in cryoSPARC v3 by patch-based motion correction, patch-based CTF correction, automated particle picking, *ab initio* reconstruction and homogeneous refinement. Two capsid maps were subsequently produced with different voxel sizes. The maps were imported into UCSF ChimeraX (*b*) and assessed for cross-correlation before the IBS micrographs were imported into Relion Image Handler and rescaled to the same pixel size as the SKKU data using the Rescale function. The data (*c*) were merged in cryoSPARC by correcting the CTF separately and merging the particles stacks at the 2D classification stage. Standard reconstruction methods were then applied as described in methods section 5 and detailed in Fig. S2 and S3. Finally, consensus maps for the capsid and tail (*d*) were obtained from the information contained within both datasets and merged to form a complete phage reconstruction. Methods adapted from Wilkinson *et al.,* **(1).**

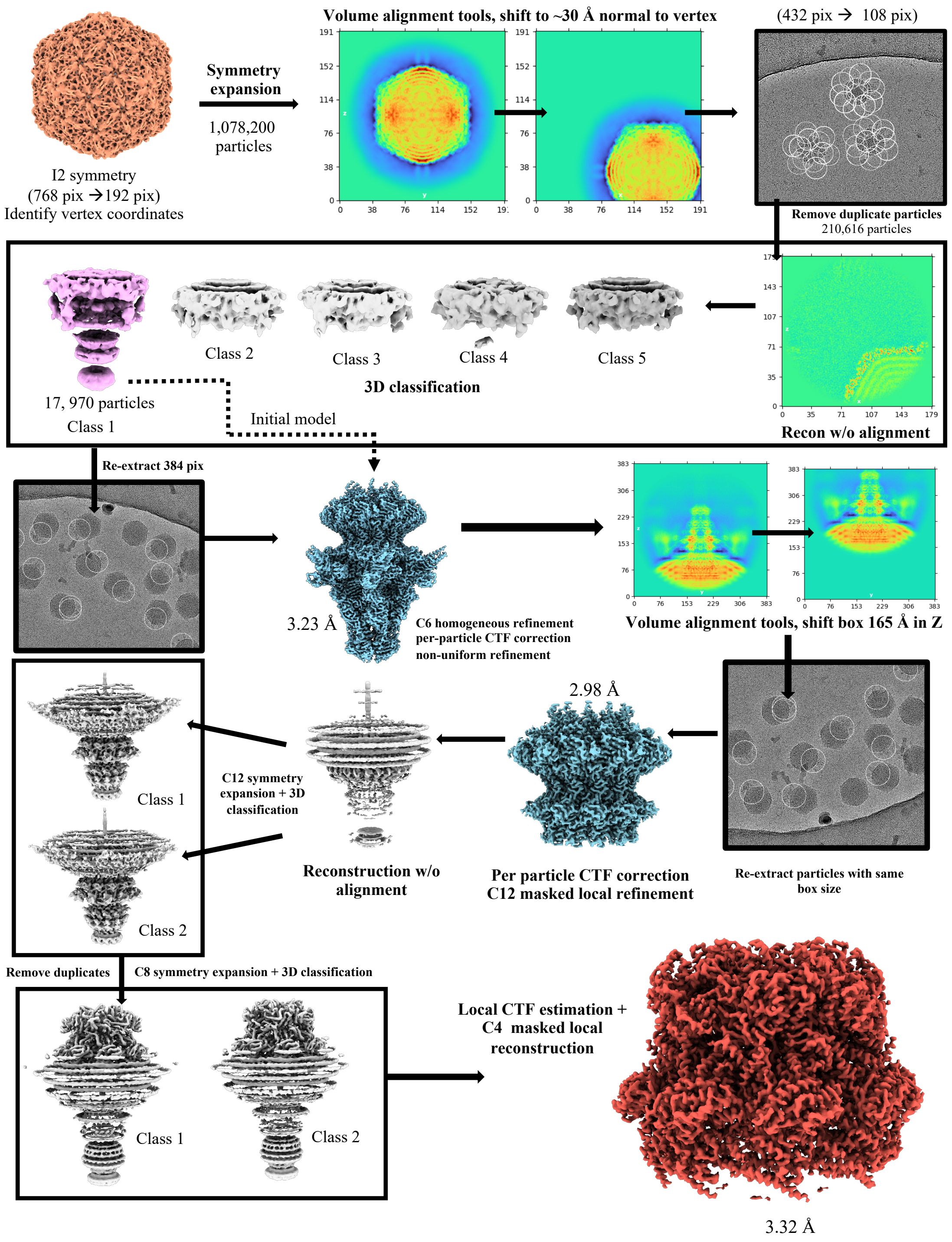


**Fig. S6. Reconstruction approach for tail and ejectosome assembly.** See supplementary method 1 for a comprehensive protocol-style description of the workflow. Reconstruction workflow was followed in cryoSPARC v4.2.1. Initial vertex coordinates were established using supplied supplementary python script (script 1: script relies on I2 capsid reconstruction). Output maps were visualized in UCSF ChimeraX v1.6.

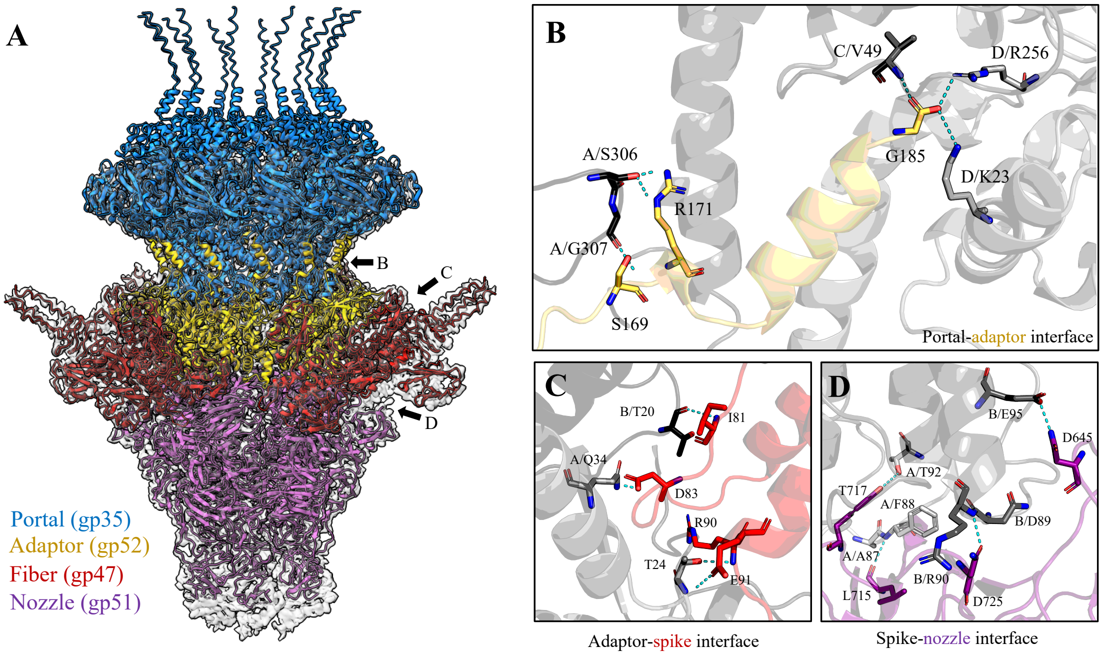


**Fig. S7. ΦM1 portal and tail machinery interfaces.** (*a*) The portal and tail machinery viewed from the side, displayed as Richardson ribbons nested within the preliminary tail density obtained at low resolution. Proteins and their gene protein identities are labelled. (*b*) The interface between the portal (shades of greys) and adaptor (yellow) involve three chains of gp35 and one chain of gp52; adaptor residue G185 interacts with two chains of gp35, while R171 and S169 interact with another chain of gp35. (*c*) The interface between the adaptor (now grey) and the spike (red) involves two adaptor chains to one spike. (*d*) The interface of the spike and nozzle.

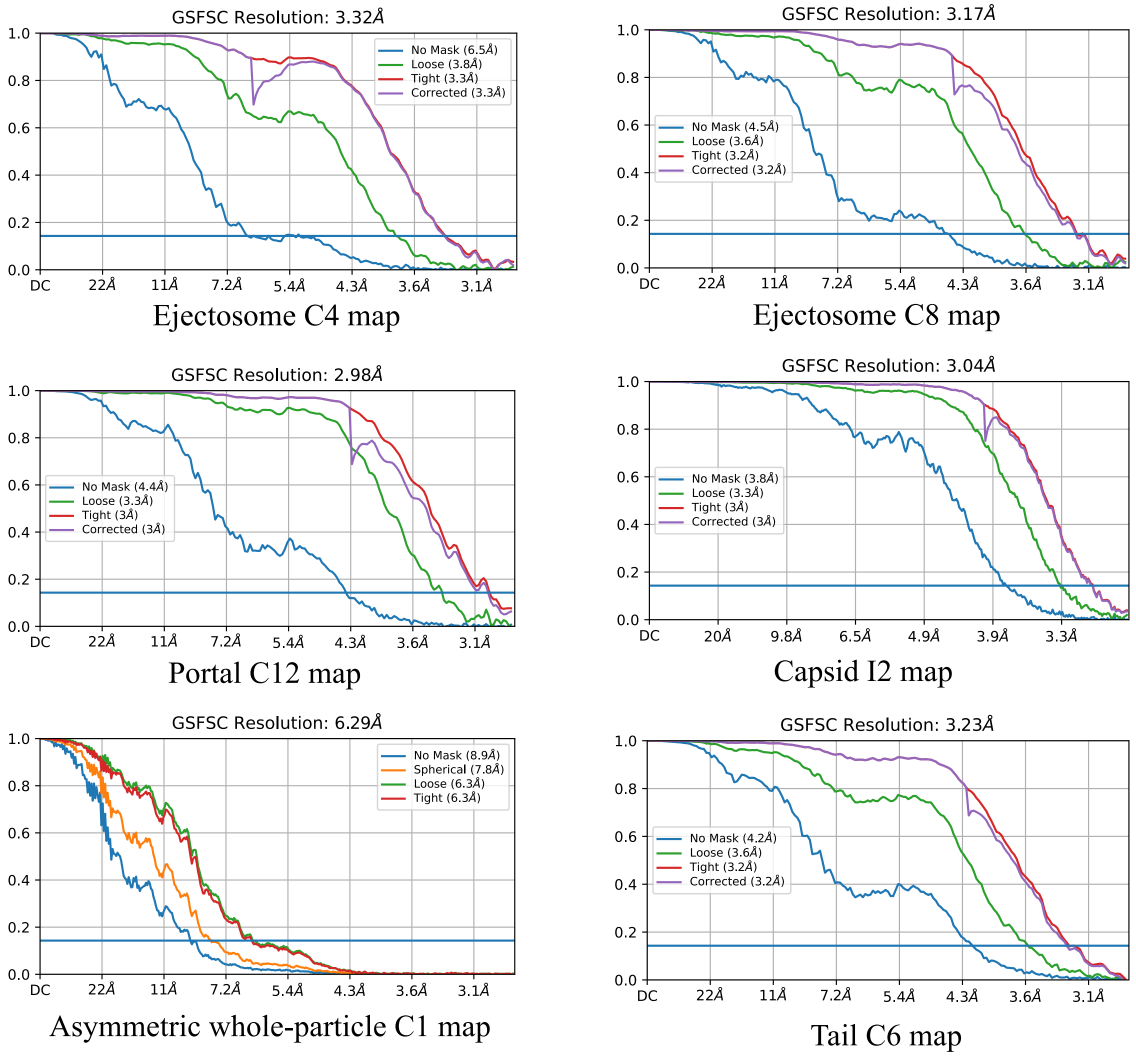


**Fig. S8. Fourier Shell Correlation Plots for *Pectobacterium* phage ΦM1 maps.** Plots were produced from non-uniform refinement jobs in cryoSPARC v4.2.1. The maps associated with each plot are listed beneath.

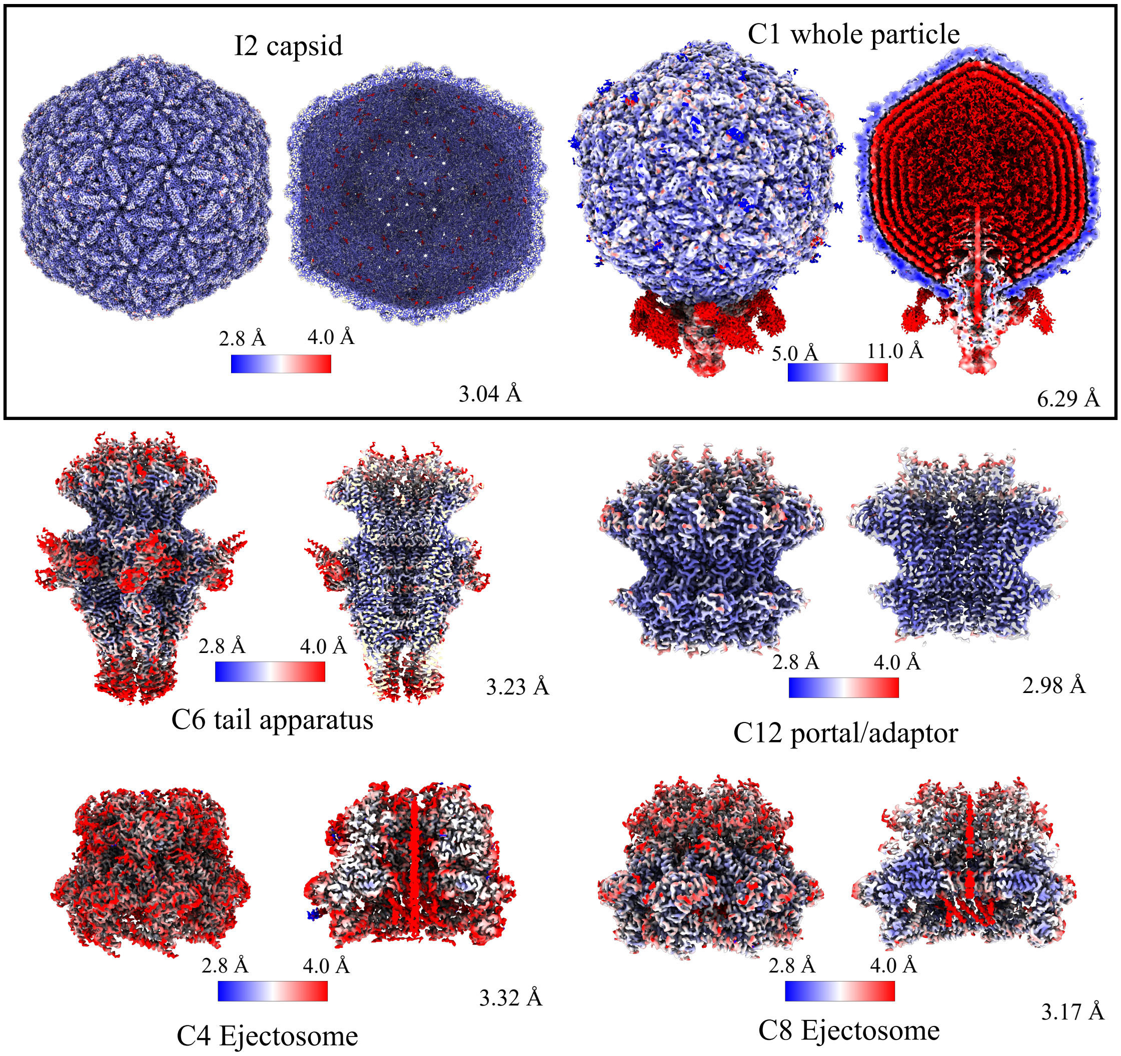


**Fig. S9. Local resolution depiction of all deposited maps.** I2 capsid contour threshold: 0.394 (PDB: 8VB0), C1 whole particle contour threshold: 0.119 (EMDB: EMD-43132), C6 spike-nozzle tail apparatus contour threshold: 0.332 (PDB: 8VBX), C12 portal-adaptor contour threshold: 0.443 (PDB: 8VB4), C4 ejectosome contour threshold: 0.205 (8VB2), C8 ejectosome contour threshold: 0.251 (EMDB: EMD-43111). Local resolution was calculated using cryoSPARC v4.2.1 Local Resolution job.

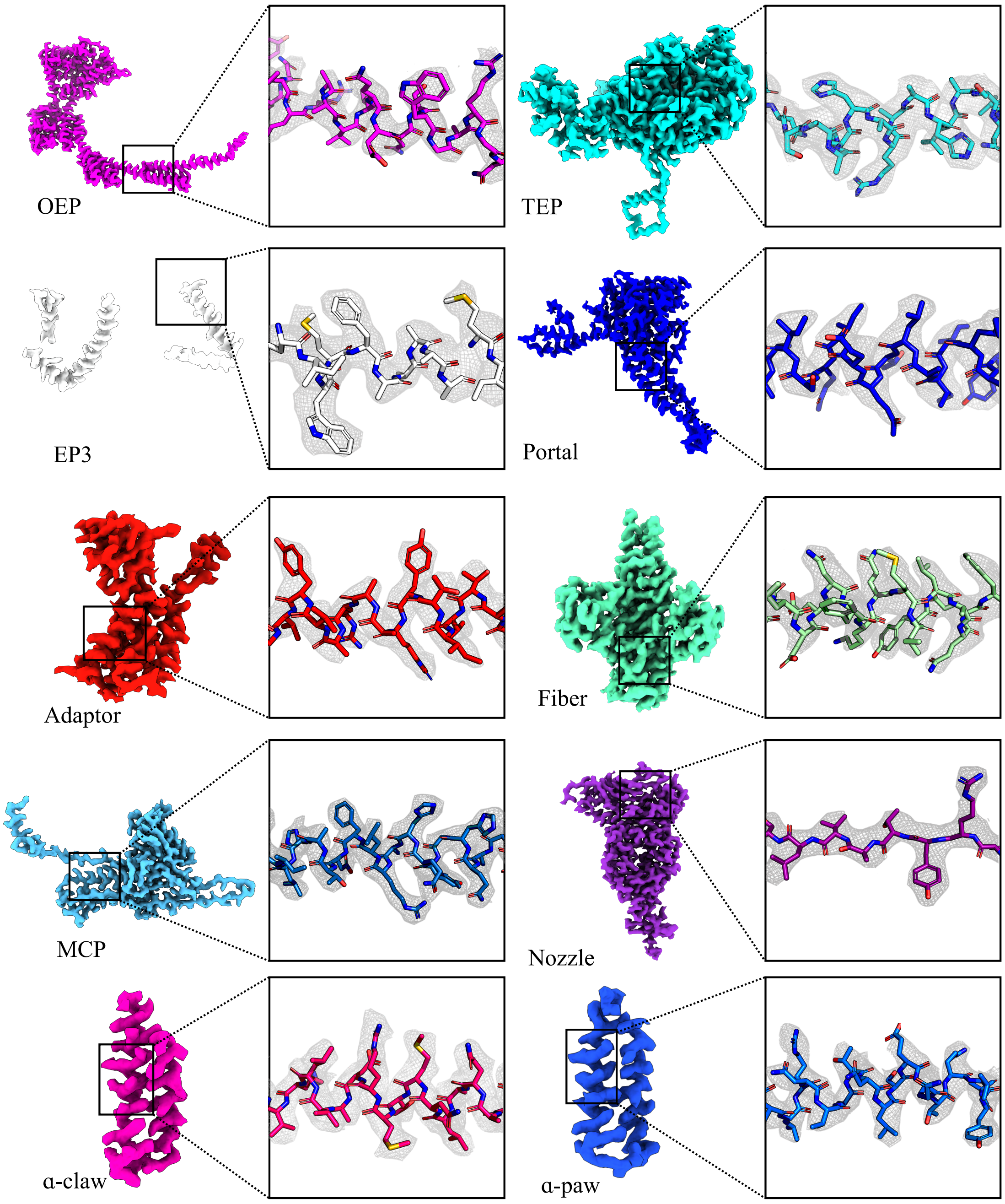


**Fig. S10. Depiction of model map quality of fit for all deposited protein models.** In each panel, left shows a zoned density corresponding to the protein (labeled below each panel). Right shows a representative helix from each model in stick representation and corresponding density. The nozzle lacks helices and so a representative beta strand is shown instead. Map contour levels are not displayed as all maps were boxed for visualization, resulting in modified density distribution.

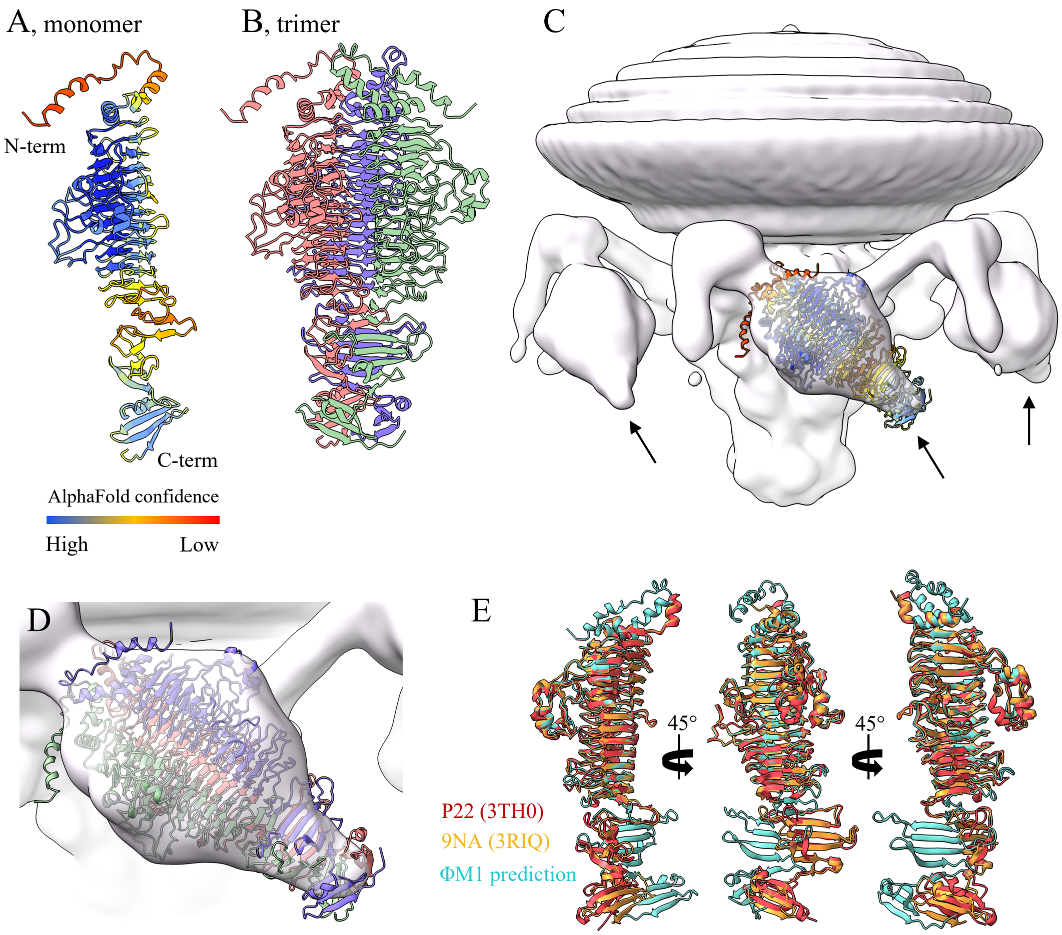


**Fig. S11. Putative globular tail spike.** The tail spike is likely encoded by gp39 of the ΦM1 genome, which is annotated in the NCBI description of the genome as 'tail spike protein'. Poor resolution of this domain precluded modelling. (*a*) An AlphaFold model of the protein (coloured by B factor) confidently predicts the core domain but loses confidence at the N- and C- termini. (*b*) The trimeric arrangement of gp39 broadly resembles the shape of the density identified for the spikes, seen in panel C marked by arrows. (*c*) A composite reconstruction of the tail spike with low-pass Gaussian filter applied. The globular domains of the tail spike density have clear 3-fold symmetric features even when reconstructed in C1, suggesting the spike is likely a trimer. The AlphaFold model coloured by B factor is fit within the filtered density with the C-terminus placed as the most distal domain. (*d*) The AlphaFold model of the gp39 is fit inside the low-pass filtered tail spike density. (*e*) A DALI structure search returned the Siphovirus 9NA tail spike receptor binding protein (PDB: 3RIQ) as the most structurally related PDB to the AlphaFold gp39 structure, followed by phage P22 tail spike (PDB: 3TH0). Overlays were produced in UCSF ChimeraX.

### SI References


1. M. E. Wilkinson, A. Kumar, A. Casañal, Methods for merging data sets in electron cryo-microscopy. *Acta Crystallogr D Struct Biol* **75**, 782-791 (2019).
2. A. Punjani, J. L. Rubinstein, D. J. Fleet, M. A. Brubaker, cryoSPARC: algorithms for rapid unsupervised cryo-EM structure determination. *Nature Methods* **14**, 290-296 (2017).
3. E. F. Pettersen *et al.*, UCSF ChimeraX: Structure visualization for researchers, educators, and developers. *Protein Sci* **30**, 70-82 (2021).
4. S. H. W. Scheres, RELION: Implementation of a Bayesian approach to cryo-EM structure determination. *Journal of Structural Biology* **180**, 519-530 (2012).
5. J. Jumper *et al.*, Highly accurate protein structure prediction with AlphaFold. *Nature* **596**, 583-589 (2021).
6. P. Emsley, B. Lohkamp, W. G. Scott, K. Cowtan, Features and development of Coot. *Acta Crystallogr D Biol Crystallogr* **66**, 486-501 (2010).
7. T. I. Croll, ISOLDE: a physically realistic environment for model building into low-resolution electron-density maps. *Acta Crystallogr D Struct Biol* **74**, 519-530 (2018).
8. M. A. Cianfrocco, M. Wong-Barnum, C. Youn, R. Wagner, A. Leschziner (2017) COSMIC2: A Science Gateway for Cryo-Electron Microscopy Structure Determination. in *Proceedings of the Practice and Experience in Advanced Research Computing 2017 on Sustainability, Success and Impact* (Association for Computing Machinery, New Orleans, LA, USA), p Article 22.
9. A. Waterhouse *et al.*, SWISS-MODEL: homology modelling of protein structures and complexes. *Nucleic Acids Res* **46**, W296-w303 (2018).
10. C. J. Williams *et al.*, MolProbity: More and better reference data for improved all-atom structure validation. *Protein Sci* **27**, 293-315 (2018).
